# Integration of genetic and epigenetic data pinpoints autoimmune specific remodelling of enhancer landscape in CD4+ T cells

**DOI:** 10.1101/2024.01.11.575022

**Authors:** Neha Daga, Nila H. Servaas, Kai Kisand, Dewi Moonen, Christian Arnold, Armando Reyes-Palomares, Epp Kaleviste, Külli Kingo, Reet Kuuse, Katrin Ulst, Lars Steinmetz, Pärt Peterson, Nikolina Nakic, Judith B. Zaugg

## Abstract

CD4+ T cells play a crucial role in adaptive immune responses and have been implicated in the pathogenesis of autoimmune diseases (ADs). Despite numerous studies, the molecular mechanisms underlying T cell dysregulation in ADs remain incompletely understood. Here, we used transcriptomic and epigenomic data from CD4+ T cells of healthy donors and patients with systemic lupus erythematosus (SLE), psoriasis, juvenile idiopathic arthritis (JIA), and Graves’ disease to investigate the role of enhancers in AD pathogenesis. By generating enhancer-based gene regulatory networks (eGRNs), we identified disease-specific dysregulated pathways and potential downstream target genes of enhancers harbouring AD-associated single-nucleotide polymorphisms, which we also validated using CRISPRi in primary CD4+ T cells. Our results suggest that alterations in the regulatory landscapes of CD4+ T cells, including enhancers, contribute to the development of ADs and provide a basis for developing new therapeutic approaches.

## Introduction

T cells play a central role in the initiation and regulation of adaptive immune responses. There are two main types of T cells: CD4+ “helper” T cells (Th) and CD8+ “cytotoxic” T cells. CD4+ T helper cells secrete cytokines that steer the immune response and activate innate immune cells, B cells, and CD8+ T cells, to respond to immunological insults. Depending on the nature of the immunological trigger, CD4+ T cells differentiate into one of several subtypes which have specialised functions in the immune response.^1^ Based on the secretion of subset specific cytokines and expression of lineage-specific transcription factors (TFs), five major Th subsets have been identified: Th1 (characterised by interferon (IFN)-γ and T-bet), Th2 (interleukins (IL)-4, IL-5, IL-13, and GATA3) T regulatory (Treg, IL-10, transforming growth factor (TGF)-β and FOXP3), Th17 (IL-17 and RORγT), and follicular T helper (Tfh, IL-21 and BCL6).^2^ Additionally, more controversial Th subsets have been described including Th9 (IL-9 and SPI1)^3^, and Th22 (IL-22 and AHR).^4^ Because of their tailored reactivity, these Th cell subsets are crucial for achieving an effective adaptive immune response.

Given their critical role in orchestrating immune responses CD4+ T cells are important players in the pathogenesis of autoimmune disease (AD): autoreactive CD4+ T cells have been implicated in systemic lupus erythematosus (SLE, a systemic AD causing widespread inflammation and organ damage)^5^, psoriasis (characterised by chronic skin inflammation)^6^, juvenile idiopathic arthritis (JIA, a rheumatic disease of childhood causing local inflammation of the joints)^7^, and Graves’ disease (mainly affecting the thyroid gland).^8^ The pathogenesis of these ADs has been linked with the dysregulation of specific Th subsets, including Tfh in SLE^9^, Th17 in psoriasis^10^, Treg in JIA^11,12^, and Th2 in Graves’s disease.^13^ Differentiation into Th subsets is driven by transcription factors (TFs), however the way in which autoreactive T cells in these diseases escape tolerance and mediate autoimmunity is incompletely understood. Genetic susceptibility along with environmental triggers likely play an important role^14^ since many ADs have common genetic predispositions, including alterations in the human leukocyte antigen (HLA) region^15^ (important for antigen presentation to T cells), as well as genes critical for T cell activation and regulation, including the IL2 receptor (CD25), CTLA4, and PTPN22.^16^ Yet, 90% of AD associated single-nucleotide polymorphisms (SNPs) lie in the non-coding part of the genome^17^, suggesting that gene regulatory elements such as enhancers play an important role in disease manifestation. In line with this, differentiation into Th subsets is driven by transcription factors (TFs) and alterations at the epigenomic level, including histone modifications at enhancers, contribute to aberrant T cell activation in ADs.^18^ However, for many AD associated (epi)genetic risk loci, we currently lack any mechanistic insights into downstream target genes and how these contribute to a loss of T cell tolerance. To address this, we have systematically integrated multiple layers of molecular genetic and epigenetic data across four types of AD.

To obtain detailed insights into the molecular mechanisms underlying T cell dysregulation in AD, and to better understand the role of enhancers in these diseases, we studied the regulatory landscapes of CD4+ T cells in SLE, psoriasis, JIA and Graves’ disease. We obtained epigenomic (H3K27ac Chromatin Immuno-preciptiation followed by sequencing - ChIP-seq) data from CD4+ T cells from these AD patients, as well as a large cohort of healthy donors, and generated enhancer-based gene regulatory networks (eGRNs), connecting TFs to enhancers harbouring AD SNPs and their putative downstream target genes. We then used these eGRNs to identify disease specific as well as general pathways that are potentially misregulated and contribute to AD pathogenesis, and used CRISPRi to validate selected disease relevant enhancer-target gene connections. Our eGRNs have the potential to uncover new genes and their upstream regulators involved in AD and may be utilized to prioritize genes for T cell-targeting treatments.

## Results

### T cells in autoimmune diseases show distinct epigenetic profiles based on histone acetylation changes between patients and controls

To investigate the role of enhancers in ADs, we performed H3K27ac ChIP-seq in CD4+ T cells obtained from peripheral blood mononuclear cells (PBMCs) of patients suffering from SLE (N=11) and psoriasis (N=15) from samples collected at initial diagnosis along with age and sex-matched healthy individuals (N=11 & N=15 for SLE and psoriasis respectively; **Figure 1A**). In addition, we obtained publicly available H3K27ac data of CD4+ T cells from patients and their respective controls suffering from Graves’ disease (N=10 patients and N=13 healthy controls; from PBMCs)^19^ and JIA (N=5; from synovial fluid (SF)).^20^ For JIA, we used T cells obtained from PBMCs from the same patients as controls (N=3), since it has previously been demonstrated that circulating CD4+ T cells from JIA patients are very similar to healthy T cells both on the epigenomic and protein level.^11,20^

**Figure 1.**
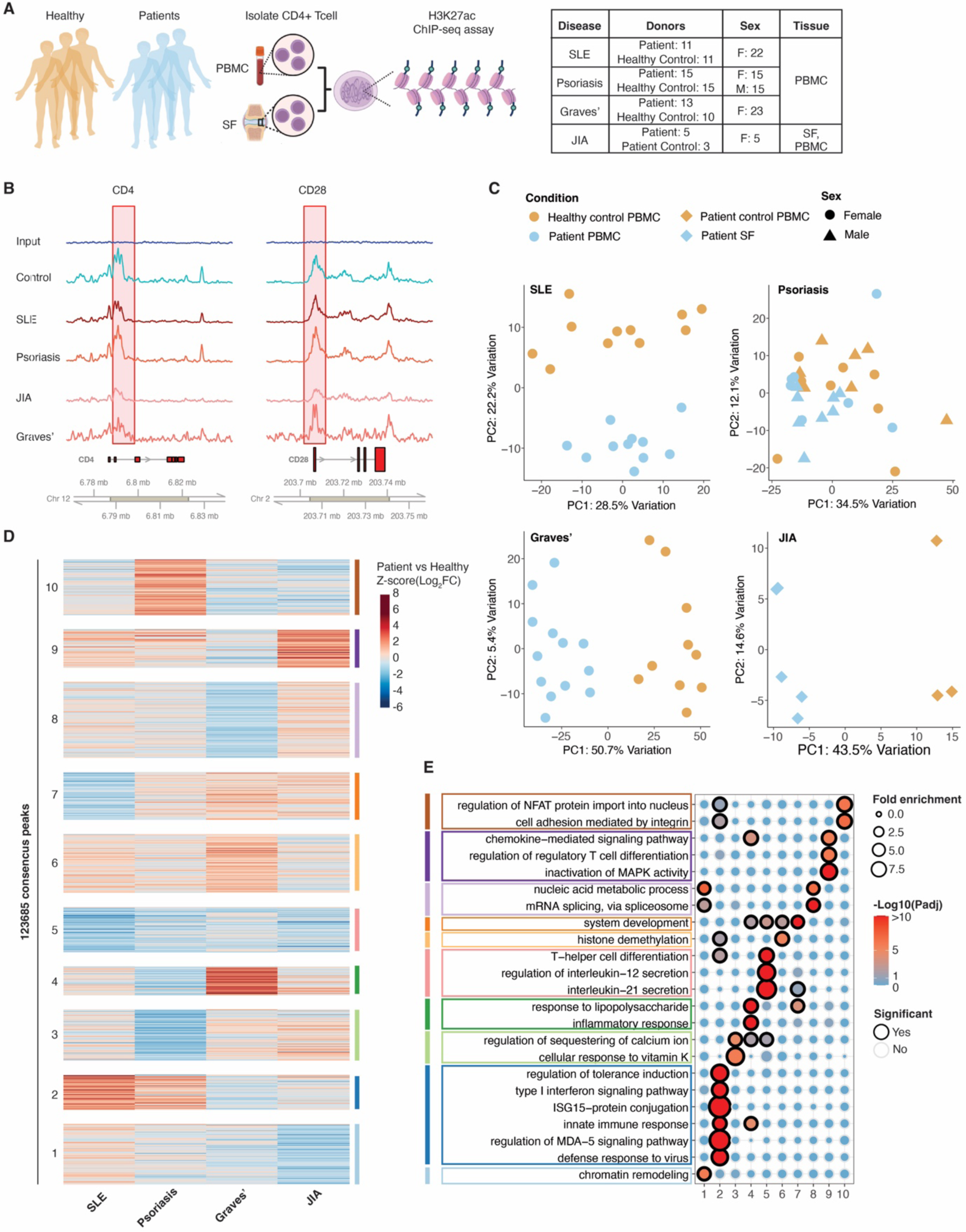
Autoimmune disease specific distinct H3K27ac profiles between patients and controls. **A)** Schematic representation of experimental procedure of CD4+ T cell isolation and H3K27ac ChIP-seq assay from patients and healthy controls; table containing sample information (F=female; M = male). **B)** Genomic tracks with H3K27ac signal around CD4 and CD28 genes for representative disease and control samples. **C)** PCA based on the top 1000 most variable acetylated peaks within each disease and corresponding control samples. **D)** K-means clustering of consensus peakset based on log2 fold changes in H3K27ac marks between patient and control samples in each disease. **E)** GO enrichment terms for each cluster in D based on closest genes using GREAT.

The newly generated H3K27ac data was of high quality and showed the expected signal around the promoter (TSS +/- 3k) and enhancer regions (**Supplemental Figure 1**), and at the CD4 (a surface receptor highly expressed in CD4+ T cells) and CD28 (an essential co-stimulatory molecule) loci, for both patient and control samples (**Figure 1B**). We called peaks in each sample using MACS2 and then generated consensus peaks per cohort (disease and control samples; see methods). This resulted in 107897, 126308, 71298 and 26730 peaks in SLE, Psoriasis, Graves’ disease and JIA respectively. Principal component analysis based on the top 1000 most variable H3K27ac peaks within each disease cohort separates patients and control groups in all diseases (**Figure 1C**), showing that the enhancer profile of T cells in AD is clearly distinct from healthy.

To compare the active chromatin regulatory landscape across the different ADs, we generated a consensus H3K27ac peak set across the four diseases, with peaks present in at least three samples, resulting in 150560 peaks. Differential acetylation analysis for each disease-control setting (using DEseq2, see methods) revealed between 5 (psoriasis) and 18k (Graves) differentially acetylated peaks (**Supplemental Figure 2 and Supplemental Table 1**). To get an unbiased view of the disease-specific enhancer profiles we grouped the variable peaks (absolute fold-change of >1 in at least one disease; 123685 peaks) into 10 clusters based on their fold-change value using k-means clustering (**Figure 1D**). While the majority of peaks are in clusters with disease-specific patterns, one cluster (cluster 5) showed uniform downregulation in all diseases. This cluster is enriched in Gene Ontology (GO) terms related to Th differentiation (**Figure 1E**; using GREAT^21^), in line with the important role of Th cells in autoimmune diseases.^22^ Cluster 9, which is upregulated in three of the four diseases, most strongly in JIA, is enriched in chemokine signalling pathways, and regulation of Treg differentiation, likely reflecting the important role of Tregs in JIA and other autoimmune diseases.^11,23^ The psoriasis-specific cluster 10 is enriched in cell adhesion processes, which is in line with disease characteristics related to disturbances in cellular adhesion that can contribute to T cell dysregulation in psoriatic skin.^24^ The Graves’ disease-specific cluster 4, is enriched in more general processes related to inflammatory response, whereas the SLE-upregulated cluster 2 is significantly enriched in processes that are important in SLE pathogenesis like type 1 IFN signaling, and defence response to virus.^25,26^ These results indicate that AD T cells are characterised by general and disease-specific enhancer signatures that potentially regulate disease relevant biological pathways.

### TF activity reveals shared and disease-specific inflammatory T cell processes across ADs

Since H3K27ac is a very dynamic histone mark^27^, it is likely that TFs are actively maintaining the AD-specific enhancer signatures. We have previously shown that changes in H3K27ac marks aggregated across TF binding sites can serve as a readout of TF activity and developed a tool (diffTF) to quantify differential TF activity.^28^ Using diffTF, we identified 341, 189, 263 and 41 TFs that are significantly differentially active between patients and controls in Graves’ disease, SLE, psoriasis and JIA, respectively (**Supplemental Table 2**), of which 100 are differentially active in at least three diseases (**Figure 2A**). Out of these, 12 TFs are commonly upregulated in patients across all diseases (**Figure 2B**). These include members of the activating protein 1 (AP-1) complex (JUN, FOS) and the NFkB complex (NFKB2, TF65), which play important roles in T cell activation, a critical process that is generally dysregulated in autoimmune conditions.^29–31^

**Figure 2.**
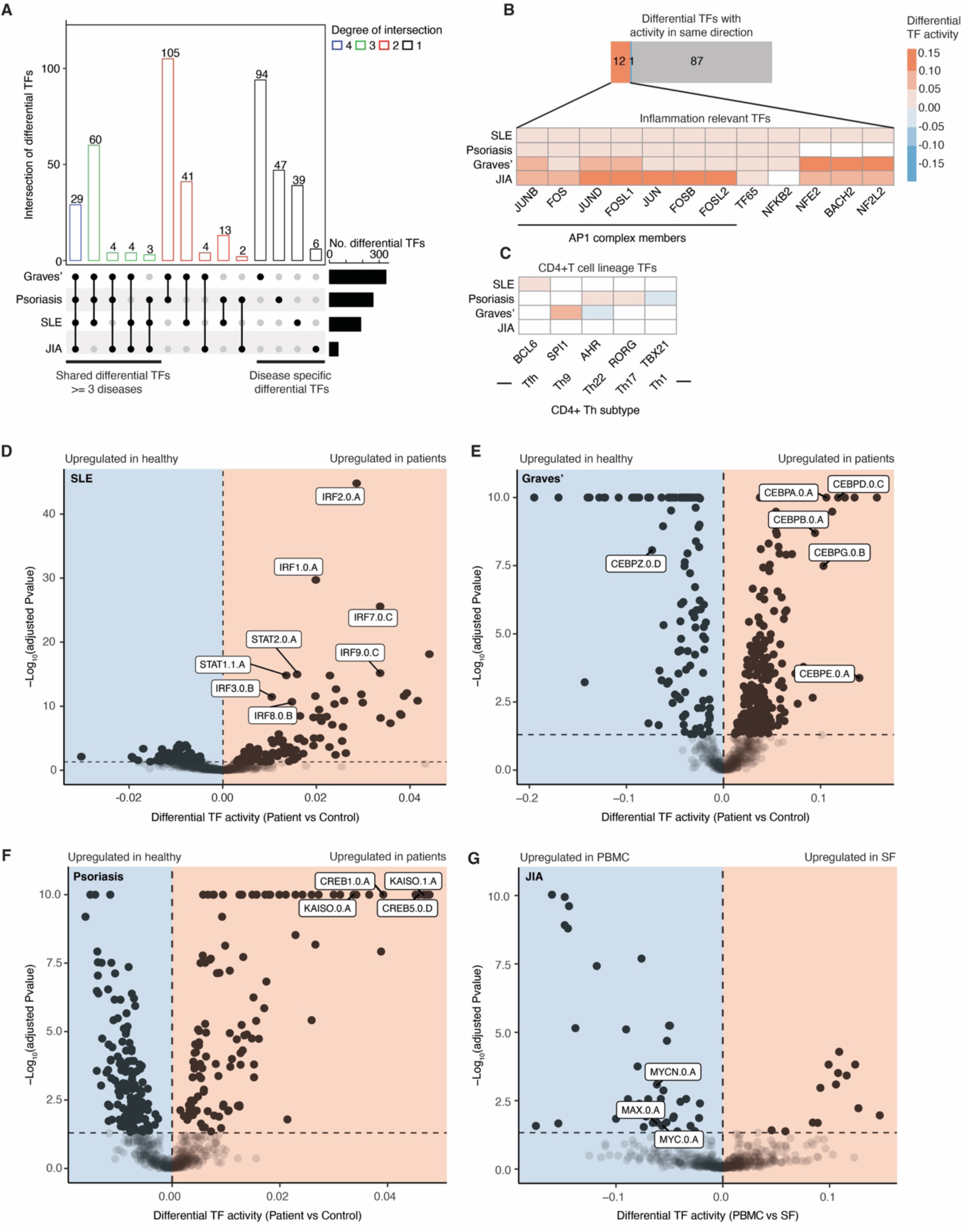
TF activity reveals shared and disease-specific inflammatory T cell processes across ADs. **A)** Upset plot of shared and unique significantly differential TFs across diseases. **B)** Inflammation relevant differential TFs (zoomed in) shared across all ADs. **C)** Disease-specific differential activity of CD4+ T helper cell lineage defining TFs. **D-G)** Volcano plot with differential TF activity (x-axis) vs log_10_ adjusted p-value (y-axis) for SLE (D), Graves’ disease (E), Psoriasis (F), and JIA (G). Selected disease specific TFs are highlighted.

We found 186 TFs are differentially active in only one of the diseases. These include several T helper cell lineage defining TFs, such as RORG (essential for Th17 differentiation^32^), AHR (essential for Th22^33^), and TBX21 (essential for Th1^34^) in psoriasis (**Figure 2C**). In SLE, we find the T follicular helper (Tfh)-specific TF BCL6^35,36^, whereas in Graves’ disease the Th9 specific TF SPI1^37^ is differentially active. These lineage specific TFs are in line with previous studies that highlighted the importance of Th17, Th22 and Th1 in psoriasis^38,39^, Tfh in SLE^9,40^, and Th9 in Graves’ disease.^41^ Other disease-specific TFs suggest further dysregulation of signalling pathways in the T cells: In SLE, we identified several IFN regulatory factors (IRF3, 5, 7, and 9) and STAT family members (STAT1, STAT2) with increased activity in patients (**Figure 2D**). These TFs are crucial for activating IFN related genes, and are involved in the aberrant expression of type I IFN genes, known as the “type I IFN signature,” an important hallmark of SLE^25,26^, and in line with the enriched GO terms in the SLE-dominated cluster 9 (**Figure 1E**). For Graves’s disease, we found high activity of the CAAT/enhancer-binding protein (C/EBP) family of TFs (**Figure 2E**), which have previously been associated with expression of inflammatory cytokines including IL-6, IL-1β, and TNF-α and regulation of immune effector functions in macrophages and neutrophils.^42^ In psoriasis, we observed a high activity of KAISO, CREB1 and CREB5 (**Figure 2F**), which are implicated in the regulation of TGFβ signalling^43–45^, a pathway that plays a major role in the pathogenesis of psoriasis.^46^ Lastly, we found that members of the Myc/Max network of TFs (MYC, MAX and NMYC) have decreased activity in T cells from JIA SF (**Figure 2G**). Myc is crucial for proper Treg function in mice, and loss of Myc in these cells contributes to development of experimental autoimmune encephalomyelitis driven by uncontrolled CD4+ and CD8+ effector T cell responses.^47^

Overall, these global analyses suggest that while ADs share a general inflammatory and T-cell activation program driven by AP-1 and NFkB, each disease also has its own distinct epigenetic profile, likely driven by a disease-specific set of TFs.

### Enhancer mediated gene regulatory network (eGRN) reveals TF-driven functional communities in CD4+ T cells

We next aimed to understand how the differentially active TFs in AD may contribute to disease mechanisms. To do so, we built an enhancer-mediated gene regulatory network (eGRN) based on paired RNA-seq and H3K27ac ChIP-seq data from PBMC-derived naive CD4+ T cells of 138 healthy donors^48^, using the eGRN inference tool GRaNIE.^49^ Briefly, GRaNIE links genes to enhancers that are then either connected to TFs or to genetic variants associated with immune-related diseases (**Figure 3A**; methods). We have previously validated GRaNIE-inferred eGRNs and demonstrated that TF-peak links recapitulate cell-type specific ChIP-seq data and peak-gene links recapitulate expression quantitative trait loci (eQTL).^49^

**Figure 3.**
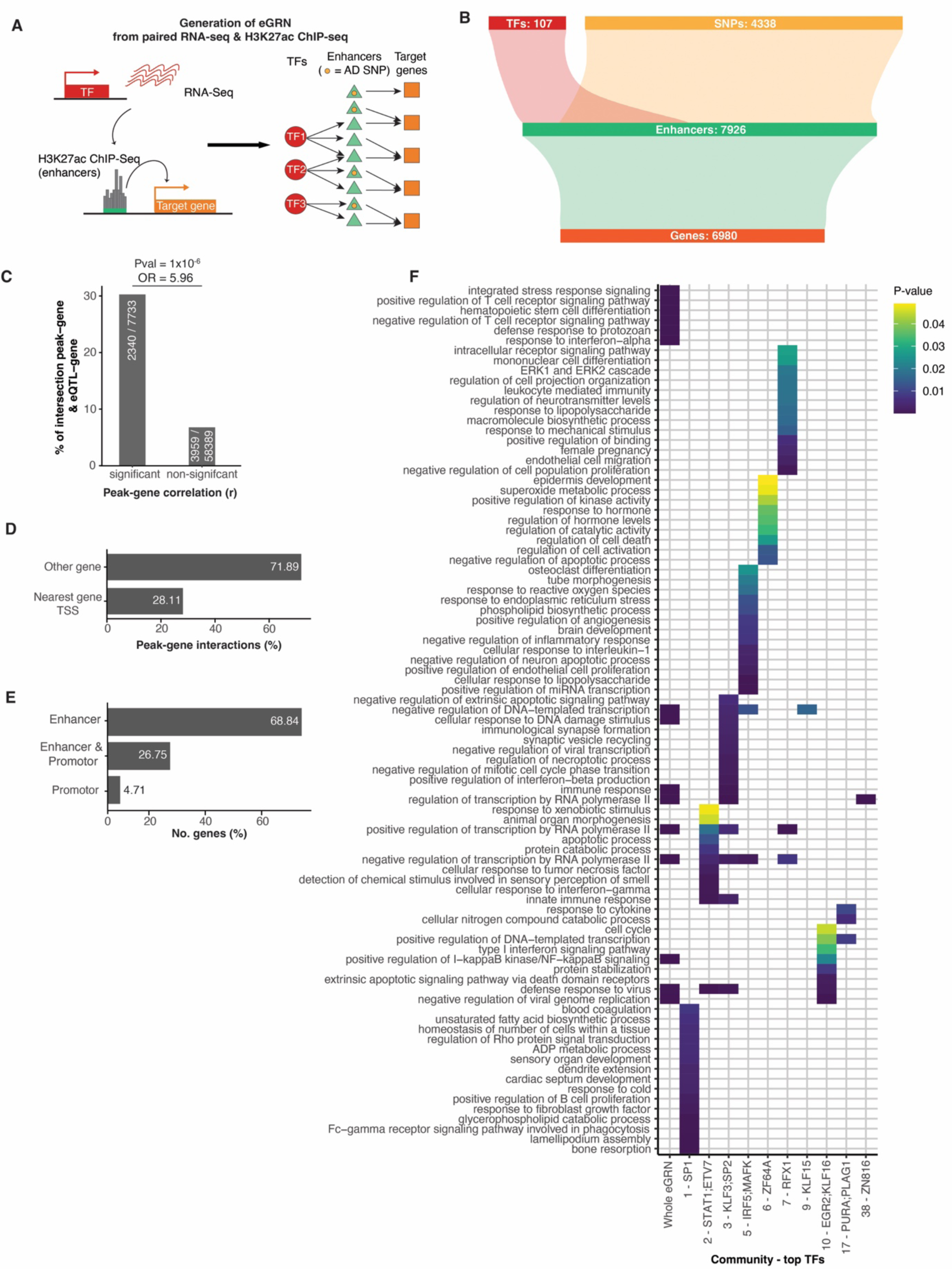
Enhancer mediated gene regulatory network (eGRN) for CD4+ T cells recapitulates known T cell biology. **A)** Schematic of T cell eGRN inference from paired RNA- and ATAC-seq data from naive CD4+ T cells from healthy individuals. **B)** Schematic representation of the naive CD4+ T cell eGRN. Numbers indicate the total number of transcription factors (TFs, red), single-nucleotide polymorphisms (SNPs, yellow), enhancers (peaks, green) and genes (orange) included in the network. **C)** Percentage of overlapping peak-gene connections from the T cell eGRN with known eQTL-gene interactions (y-axis). Data is shown for significant and non-significant peak-gene correlation pairs (x-axis) obtained from the naive CD4+ T cell eGRN. White numbers in bars indicate the number of overlapping genes / total number of genes. **D)** Percentage of peak-gene interactions (x-axis) from the T cell eGRN with peaks associated to the transcription start site (TSS) of the nearest genes, or another gene (y-axis). **E)** Percentage of genes (x-axis) from the naive CD4+ T cell eGRN linked to enhancers, enhancers & promoters or only promoters (y-axis). **F)** GO enrichment of biological processes based on all genes included in the whole naive CD4+ T cell eGRN (whole eGRN) and specific communities (indicated by numbers, x-axis). Top TFs for each community (based on degree) are highlighted.

The resulting CD4+ T cell eGRN comprises 174 TFs (of which 80 were also differentially active in at least one of the ADs), and 4338 AD associated GWAS variants connected to 6980 genes through 7926 unique enhancers (**Figure 3B, Supplemental Table 3**). We validated the network by overlapping T cell specific expression quantitative trait loci (eQTLs)^50^ with enhancer-gene pairs in the eGRN and found them strongly enriched over non-significant pairs (OR>5; p-value<1e-6; **Figure 3C**). Notably, 78% of enhancers in the eGRN are not connected to their nearest gene (**Figure 3D**), which is in line with previous observations^51^, and highlights the importance of eGRNs to identify the putative target gene of an enhancer. Categorising, the peak-gene links as either promoter (+/- 2kb TSS) or enhancer (>2kb from TSS), we find that 70% of the genes in the eGRN are linked to enhancers, highlighting the importance of considering distal regulatory regions (**Figure 3E**).

GO term analysis of all genes included in the eGRN showed an enrichment in pathways highly relevant for T cell biology, including ‘immune response’, ‘regulation of T cell receptor signalling’, ‘regulation of NFkB signalling’ and ‘type I IFN signalling pathway’ (**Figure 3F**, whole eGRN). When we cluster the eGRN into communities using louvain based clustering, we find 48 communities (**Supplemental Table 4)**, each regulated by a small set of TFs and enriched for distinct biological processes (**Figure 3F, Supplemental Figure 3**). Out of these communities, several contain TFs that are differentially active across ADs. For example, community 2, enriched for immunological synapse formation and cellular response to IFN-γ and TNF, is regulated by STAT1 and ETV7 (the TFs with the highest degree in this community, **Supplemental Figure 3**) that also have an increased activity in SLE (**Figure 3D**). Community 7, enriched for response to stimulus and leukocyte mediated immunity is mainly regulated by RFX1, and also comprises the AP-1 complex TFs JUND, FOS, FOSB and FOSL1 that showed increased activity in all ADs (**Figure 2B**). Community 5 contained CEBPB which has a disease specific increased activity in Graves’ disease (**Figure 2E**), and community 33 contained MYC which has a disease specific decreased activity in JIA SF (**Figure 2G**), though these were not significantly enriched for any biological processes.

In summary, the CD4+ T cell specific eGRN comprising enhancers, TFs and genes, captures relevant biological processes related to T cell function relevant for ADs, and many communities are regulated by TFs that are differentially active in AD. Thus, the naive T cell eGRN from healthy donors captures AD signatures, and can be used to identify disease specific TFs, regulatory elements and their downstream effector genes.

### Disease specific eGRNs integrating genomic and epigenomic data capture potential dysregulated pathways in AD T cells

We next used the CD4+ T cell eGRN to integrate disease-specific genetic (GWAS) and epigenetic (differential TFs and enhancers) evidence to understand T cell dysregulation in AD. To do so, we annotated the eGRN nodes (TFs and H3K27ac peaks) with four types of molecular disease-evidence: (i) differential TF activity between autoimmune patients (SLE, psoriasis, Graves’ disease and JIA) and controls, (ii) H3K27ac peaks that overlap with differential H3K27ac signal between autoimmune patients and controls, (iii) H3K27ac peaks that overlap with SNPs associated to AD from the NHGRI-EBI GWAS catalogue^52^, and (iv) H3K27ac peaks that overlap with peaks associated with significant colocalized histone quantitative trait loci (hQTLs)-AD GWAS SNPs.^53^ Out of these, i and ii reflect epigenetic disease evidence, while iii and iv reflect genetic disease evidence.

We subsetted the CD4+ T cell eGRN into four disease specific eGRNs (for SLE, psoriasis, Graves’ disease, and JIA) by retaining gene links connected to at least two upstream nodes (TF or enhancer) with molecular disease evidence (schematic in **Figure 4A**). Additionally, we constructed one general AD eGRN comprising genes connected to upstream nodes (TFs, Peaks) associated with molecular disease evidence from at least two diseases. This resulted in five different AD eGRNs (**Figure 4B, Supplemental Table 5)**. Genes from these AD eGRNs exhibit some degree of overlap between two or more networks/diseases, suggesting these diseases share common disease pathways (**Figure 4C**).

**Figure 4.**
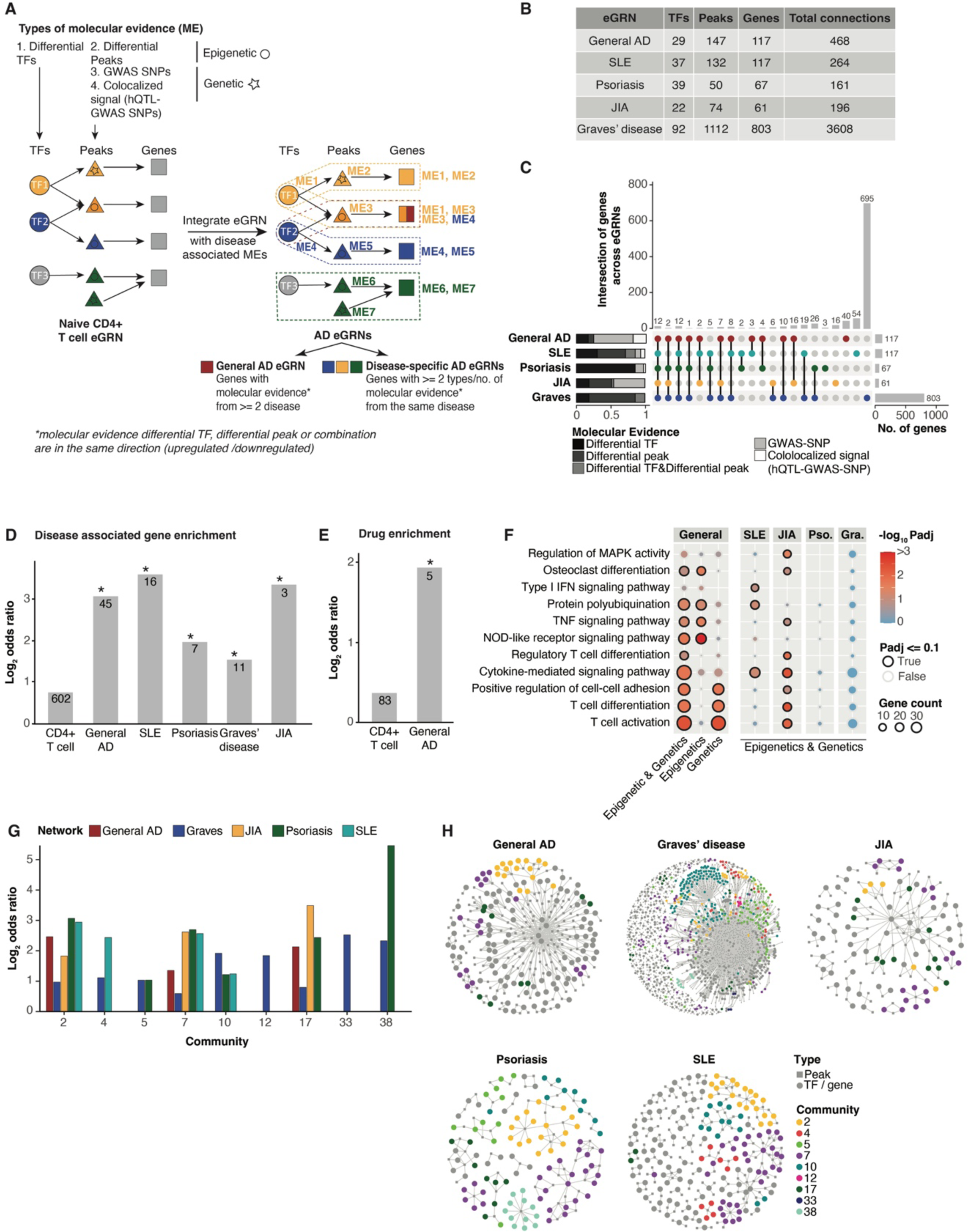
AD specific eGRNs integrating genomic and epigenomic data are enriched for disease associated genes, drug targets and relevant biological pathways. **A)** Overview of AD eGRNs generation method from the CD4+ T cell eGRN through overlap with disease specific molecular evidence (ME). **B)** Number of unique transcription factors (TFs), peaks, genes and total connections included in each disease specific eGRN. **C)** Upset plot showing the intersection of genes across the disease specific eGRNs. Shades in barplots indicate the proportion of genes with specific ME. **D-E)** Enrichment of genes from the disease specific eGRNs (x-axis) with disease associated genes (D) and known drug targets obtained from the Open Targets Platform^54^ (D). Number of genes are indicated in the bars (*= Fisher’s exact test p-value <0.05 & odds ratio > 1). **F)** GO term enrichment analysis of AD eGRNs (columns). Pso. = psoriasis, Gra. = Graves’ disease. **G)** Enrichment of community genes from Figure 3F (x-axis) across AD eGRNs. Only significant enrichments are shown (Fisher’s exact test p-value < 0.05 & odds ratio > 1). **H)** Force-directed visualisation of the AD eGRNs. Colours correspond to genes coming belonging to enriched communities.

We overlapped the eGRN genes with known disease-associated genes from the OpenTargets platform.^54^ This revealed a significant enrichment among genes from the respective disease specific networks (SLE, psoriasis, Graves’ disease and JIA), while the general AD eGRN was significantly enriched in disease genes associated with any AD (**Figure 4D**). Since disease-gene annotations are partially derived from GWAS studies, which may create a circular argument for the part of our network that is based on genetic evidence, we confirmed that a similar enrichment was observed when considering AD eGRNs without genetic evidence (**Supplemental Figure 4**). Furthermore, the general AD eGRN is enriched for genes that are known drug targets for ADs (**Figure 4E**).

We next asked whether genetic and epigenetic disease evidence converges onto the same biological pathways. We performed GO term enrichment analysis for genes in the general AD eGRN with only genetic evidence (GWAS-SNPs & co-localised signal between hQTLs and GWAS-SNPs) and for those with only epigenetic evidence (differential TFs & differential peaks). This revealed complementary processes enriched in these two subsets: genes in the epigenetic subset enriched in immune signalling related pathways (i.e. TNF and NOD-like receptor signalling), while genes in the genetic subset were enriched in processes related to T cell differentiation and activation (**Figure 4F**). Notably, when combining genetic and epigenetic evidence, additional relevant processes like cytokine-mediated signaling and Treg differentiation were enriched. This highlights the power of combining genetic and epigenetic disease evidence in eGRNs for understanding disease mechanisms.

The general AD eGRN captures processes relevant for T cell activation and differentiation (important for AD pathogenesis), whereas genes in the disease-specific AD eGRNs are enriched in biological processes that are important for disease specific pathogenesis (**Figure 4F**). For instance, genes in the SLE network are significantly enriched in type 1 IFN related signalling, an important pathway known to be mis-regulated in SLE.^25,26^ We observed genes in the JIA network to be enriched in osteoclast differentiation, a process related to synovial joint inflammation characterising JIA patients, as well as regulation of Treg differentiation, a known cell type involved in JIA pathogenesis.^11,20^

Lastly, we checked whether the AD eGRNs were enriched in genes from specific communities which were identified in the naive CD4+ T cell eGRN (**Figure 3F**). We found a significant enrichment of community 2 and 7 across all AD eGRNs, and community 17 across the general AD, Graves’ disease, JIA and psoriasis eGRNs (**Figure 4G/H**). All of these communities are enriched in pathway genes related to immune activation (**Figure 3F**) and included TFs that are highly relevant for T cell activation (including AP-1 complex members in community 7, STAT1 in community 2 and PURA in community 17)^31,55^, again highlighting shared mechanisms of T cell activation across different ADs. We also identified more specific enrichments of communities across the different AD eGRNs, including community 4 (most enriched in SLE), community 38 (most enriched in psoriasis), and communities 12 and 33 (only enriched in Graves’ disease) (**Figure 4G/H**). These results further show that our AD eGRNs recapitulate disease relevant pathways, with shared as well as distinct gene signatures.

### Identification of Th subset gene signatures linked to specific AD eGRNs

T cell activation state and differentiation into distinct Th subsets play an important role in ADs.^22^ Therefore, we asked whether the general AD and disease-specific eGRNs are enriched for genes related to distinct Th subsets. We obtained RNA-seq data from five different CD4+ T cell subtypes from healthy donors (Th1, Th2, Th17, Treg and Tfh) that were stimulated with anti CD3/CD28 beads^56^, and defined stimulation-responsive genes as those that are significantly differentially expressed in comparison to resting cells using DESeq2 (p-adjusted<= 0.1, see methods). All AD eGRNs were significantly enriched in stimulation-responsive genes from one or more CD4+ T cell subtypes (**Figure 5A, Supplemental Table 6**). The strongest enrichments were observed for Th1, Th17, and Tregs in the psoriasis eGRN, and Tregs in the JIA eGRN, in line with the cell types important in these diseases.^11,20,57^ Stimulation-responsive genes in Tregs were also significantly enriched in the Graves’ disease and general AD eGRNs (although less strong than in JIA), likely reflecting the crucial role of Tregs in maintaining general immune tolerance and in line with their suggested dysregulation in ADs in general.^58^

**Figure 5.**
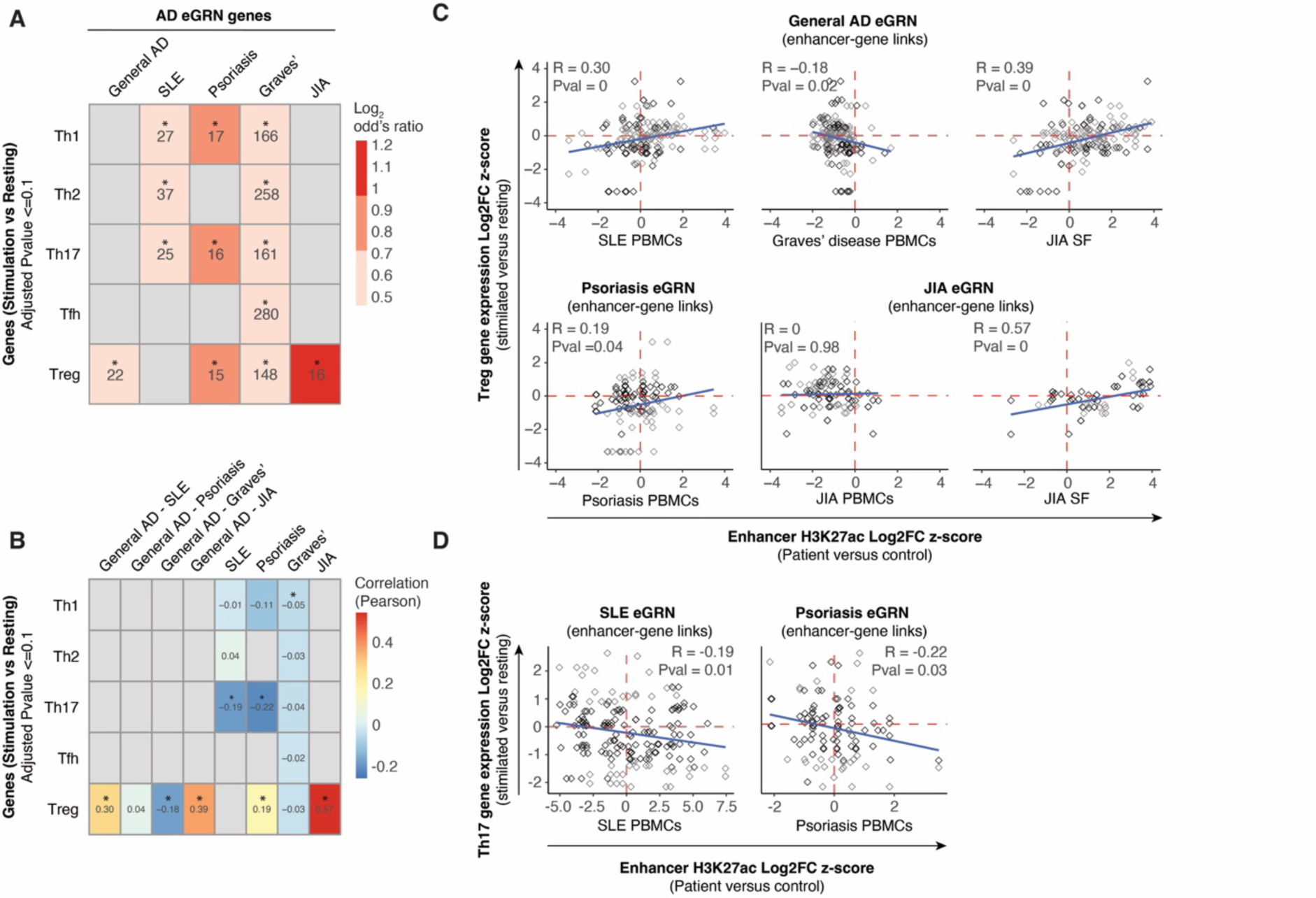
AD specific eGRNs are enriched in regulatory elements driving gene expression related to T helper subset differentiation. **A)** Enrichment (log2 odds ratio, colour scale) of T helper (Th) subset specific stimulation associated genes (rows) in the AD eGRNs (columns). Numbers in cells indicate the number of overlapping genes (* = Fishers’ exact test p-value <0.05; grey cells are not significant. **B)** Pearson correlation (colour scale) between expression of Th subset specific stimulation associated genes (rows) and differential acetylation of peaks from the AD eGRNs (columns). *= Pearson correlation p-value <0.05. **C)** Pearson correlation of gene expression changes in stimulated versus resting Tregs (y-axis) with H3K27ac changes in autoimmune patients versus controls (x-axis). Titles indicate eGRN from which enhancer-gene links were taken. Blue lines: regression lines. Genes that change significantly in Tregs upon stimulation are highlighted in grey (adjusted p-value <0.1). **D)** Correlation of gene expression changes in stimulated versus resting Th17 (y-axis) with H3K27ac changes in autoimmune patients versus controls (x-axis).

Next, we aimed to understand how enhancer remodelling in AD patients contributes to Th subset-specific activation. We used the AD eGRNs to link enhancers to stimulation responsive genes, and assessed the correlation between disease-specific changes in H3K27ac signal (AD patients versus controls) and differential expression of the connected gene upon stimulation of the Th subset (**Figure 5B**). We restricted this analysis to the Th-subsets that were enriched in the respective eGRN. We found a positive correlation between expression changes in Tregs with enhancer activation in JIA and SLE, and a negative correlation with enhancers in Graves’ disease (**Figure 5B/C**). This may reflect the distinct roles that Tregs plays in these diseases and is in line with reports of Treg down-regulation in Graves’ disease^59^, and their upregulation in JIA SF.^11^ For JIA, the same positive correlation was observed for the JIA-specific eGRN for enhancer activity in synovial fluid, but not for enhancer activity in PBMCs (**Figure 5C**, bottom right panels), in line with the local nature of this disease.

For psoriasis H3K27ac changes in patients were positively correlated with gene expression changes in stimulated Tregs and negatively correlated with expression changes upon stimulation of Th17 (also enriched among the psoriasis eGRN; **Figure 5C/D**). Th17 has been recognized as a crucial T cell subset driving psoriasis pathogenesis, and it has been suggested that Tregs in psoriasis can differentiate into IL-17-producing cells that are different from “classical” Th17^60^, in line with what we observe in our correlation analysis.

The SLE-specific eGRN revealed a negative correlation of SLE enhancer activity and activation response in Th17 (**Figure 5D**). Many genes that have increased enhancer activity in SLE, and decreased expression in Th17, are related to IFN responses (i.e. IFI44, OAS3, MX2, **Supplemental Table 6**). It has previously been shown that, under inflammatory conditions, Th17 cells can start producing IFN-γ in addition to their classical IL-17 cytokine profile, and contribute to autoimmunity.^61^ Notably, levels of both IFN-γ and IL-17 are increased in SLE patients^62^, suggesting a potential role for non-classical IFN-γ producing Th17 cells in the disease pathogenesis.

Overall, we found disease-specific T cell activation patterns in (non-classical) T cell subtypes reflected in the disease-specific enhancer remodelling.

### JIA eGRN enhancer-genes within SEs respond to BET inhibitor JQ1 revealing disease relevant network connections

Super-enhancers (SEs) are stretches or clusters of enhancers in close proximity that are enriched in active enhancer marks (H3K27ac) and TFs.^63^ They drive robust gene expression at a much higher level than individual enhancers, and regulate genes involved in cell identity, state and development. SEs in CD4+ T cells are reportedly enriched for AD GWAS-SNPs.^17^ Here we used our eGRN framework to investigate the mechanisms underlying this enrichment. To this end, we obtained SE annotations for resting and stimulated naive and memory CD4+ T cells^64^, and overlaid them with our AD eGRNs. We observed that enhancers in the JIA, psoriasis and general AD eGRNs are significantly enriched in T cell specific SEs (**Figure 6A**). Furthermore, eGRN enhancers located within these SEs were significantly more acetylated in patients than controls when compared to enhancers not overlapping SEs across all eGRNs except for Graves’ disease (**Figure 6B**). These results highlight the relevance of CD4+ T-cell associated SEs in AD-specific epigenetic remodelling.

**Figure 6.**
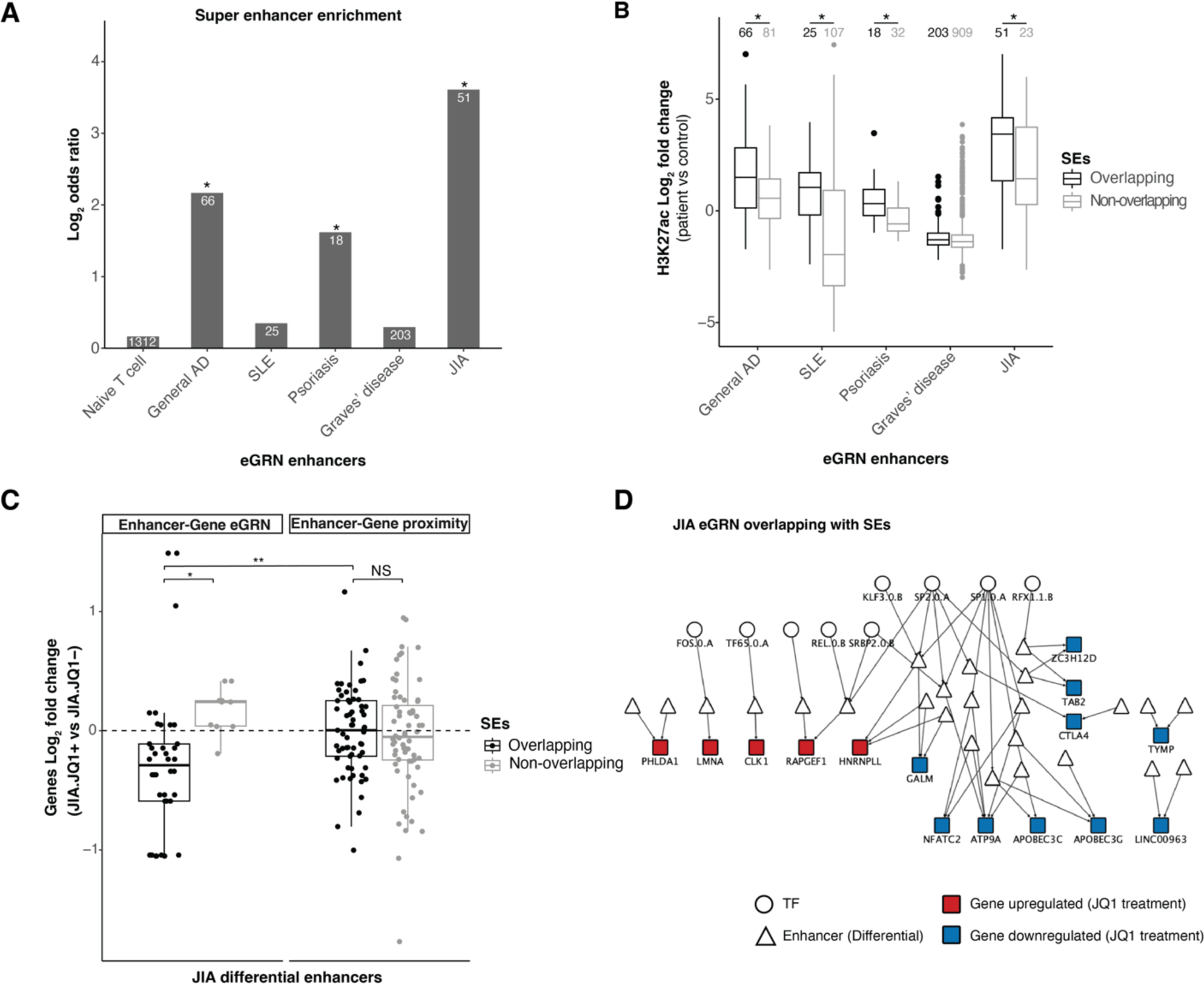
AD eGRNs are significantly enriched in Super Enhancers (SEs) and genes connected to SEs in JIA eGRN respond to BET inhibitor treatment. **A)** Enrichment (log2 odds ratio, y-axis) of T cell specific SEs across the naive CD4+ T cell eGRN and AD specific eGRNs (x-axis) (Fishers’ exact test, * = p-value <0.05). Number of genes from the eGRNs overlapping with super enhancers are indicated. **B)** H3K27ac changes in autoimmune patients versus controls (y-axis) for enhancers from AD specific eGRNs. Enhancers are separated based on their overlap with T cell specific super enhancers (black = overlapping, grey = non-overlapping). Number of unique enhancers is indicated. * = p-value <0.05 (T test). **C)** Gene expression changes in JIA patients upon BET inhibitor treatment versus no treatment (JQ1+ vs JQ1-, y-axis) for genes connected to JIA-specific differential enhancers from JIA-eGRN (Enhancer-Gene eGRN) and enhancer-gene connected only by proximity (Enhancer-Gene proximity). Genes are stratified based on whether they are connected to T cell specific SE (black = overlapping, grey = non-overlapping) * = p-value <=0.1; ** = p-value <=0.01; NS = Non-significant (T test). **D)** JIA-eGRN with TF, enhancer and gene connection for JIA specific differential enhancers within SEs in JIA-eGRN

Next, we used publicly available RNA-seq data from synovium-derived CD4+ T cells from JIA patients before and after treatment with the BET inhibitor JQ1, known to target SEs.^20^ Using enhancer-gene links from the JIA eGRN, we observed that genes connected to SEs overlapping differential enhancers in JIA were more down-regulated upon JQ1 treatment than non-SE connected genes within the JIA eGRN (p-value 0.1), while SE inhibition had no effect on genes not in the JIA eGRN, suggesting JQ1 acts exclusively on SEs that are remodelled in JIA (**Figure 6C**). Notably, genes in the JIA eGRN were more strongly down-regulated than genes linked to SEs by proximity (+/ 2kb, **Figure 6C**), with a p-value of 0.008 and a p-value of 0.25 (limited to only unique genes connected to enhancers). These observations suggest that the JQ1 treatment (which affects super enhancers) seems to act by targeting the enhancers and thus affecting their downstream target gene connections in the JIA eGRN. This provides orthogonal validation of the disease-relevance of the enhancer-target-gene links identified through our integrative approach of building eGRNs with disease specific epigenetic and genetic information.

These JIA eGRN genes connected to SEs (**Figure 6D**) include CTLA4 and NFAT4 genes known to be important in the T-cell activation processes^65–67^, which show down-regulation upon JQ1 treatment.

### Prioritisation and validation of eGRN enhancers-gene pairs potentially important for T cell dysregulation in AD

To identify the top shared genes that are potentially mis-regulated across several ADs, we ranked the genes based on i) the number of AD eGRNs they are part of ii) number of molecular disease evidence types and iii) total number of molecular evidences (**Figure 7A**). To add more information to the prioritised genes, we used three additional metrics i) regulation by a fine mapped autoimmune disease variant ii) significant change in gene expression upon T cell stimulation iii) annotation from known association to autoimmune condition.^54^ The top three candidates in our prioritised gene list are *IKZF3*, *CTLA4* and *CDK12*, which have previously been identified as crucial regulators of T cell activation.^65,68,69^ One of the enhancers connected to *CTLA4* (chr2:203930706-203942262), was also connected to *ICOS* and *CD28* in the T cell eGRN (**Figure 7B**). These three genes, all members of the immunoglobulin supergene family, play important distinct as well as synergistic roles in the regulation of T cell responses^66^, and we observed that their expression was significantly upregulated in all subsets of activated T cells (**Figure 7C**). Enhancer chr2:203930706-203942262 (**Figure 7B**, highlighted in grey) overlaps two differential AD associated peaks, both of which were downregulated in Graves’ disease (**Figure 7D**), and one upregulated in JIA (**Figure 7D**). TF activity of KLF3 and SP2, which have binding motifs on this enhancer, was significantly reduced in Graves’ disease (**Figure 7E**). To validate the enhancer-gene links, we performed CRISPR interference (CRISPRi) in activated primary human T cells using four single-guide RNAs (sgRNAs) targeting enhancer chr2:203930706-203942262 (sgRNA design can be found in **Supplemental Figure 5 and Supplemental Table 7**). Targeting the enhancer led to an overall decrease in *ICOS*, *CTLA4* and *CD28* expression (**Figure 7F**). The observed downregulation was strongest in *ICOS*, followed by *CTLA4* and *CD28*, in line with their increased genomic distance from the perturbed enhancer (**Figure 7B**).

**Figure 7.**
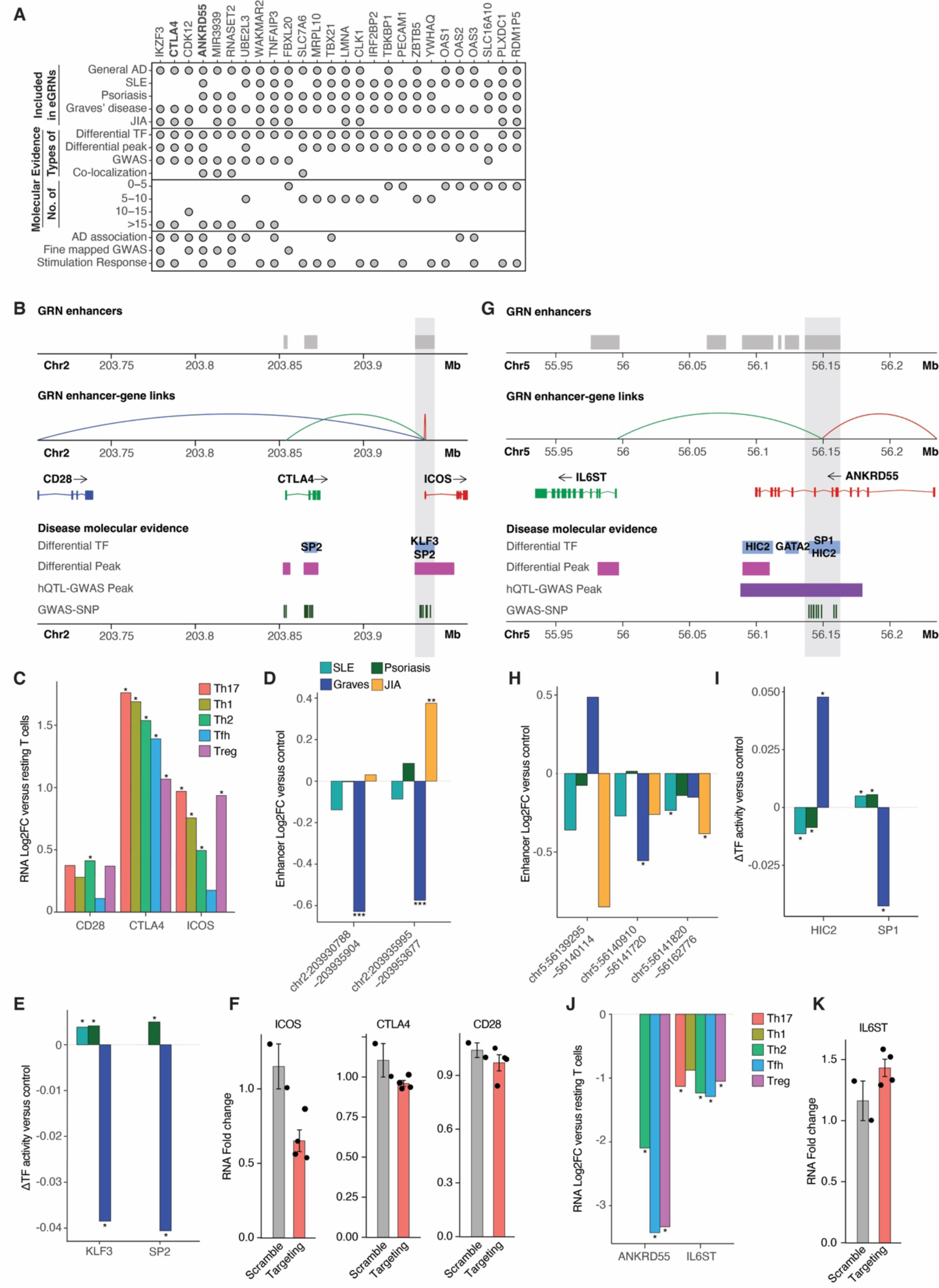
Prioritisation and validation of disease relevant enhancers from the AD eGRNs. **A**) Ranked list of prioritised genes relevant for AD (y-axis) based on their presence in AD specific eGRNs, and different types of molecular evidence associated with them (x-axis). **B**) Genomic context of *ICOS*/*CTLA4*/*CD28* and their links to AD eGRN enhancers (top grey boxes), binding sites for differential TF (blue boxes with TFs highlighted in black), differential H3K27ac peaks in AD (pink boxes), hQTL-GWAS peaks (purple boxes) and GWAS-SNPs associated with AD (dark green stripes). **C**) Change in gene expression of *ICOS*/*CTLA4*/*CD28* in stimulated versus resting CD4+ T cells (DESeq2 log2 fold change, y-axis). **D**) Change in H3K27ac signal of enhancers from the AD eGRNs linked to *ICOS*/*CTLA4*/*CD28* in AD patients versus controls (DESeq2 log2 fold change, y-axis). **E**) Differential TF activity in AD patients versus controls (calculated using DiffTF, y-axis) for KLF3 and SP2. **F)** Expression of *ICOS*/*CTLA4*/*CD28* upon CRISPRi-mediated enhancer inhibition in primary human CD4+ T cells. mRNA expression relative to one non-targeting control gRNA (scramble) is shown as mean ± SEM. **G**) Genomic context of *ANKRD55*/*IL6ST* and their links to AD eGRN enhancers. **H**) Change in H3K27ac signal of enhancers from the AD eGRNs linked to *ANKRD55*/*IL6ST* (x-axis). **I**) Differential TF activity in AD patients versus controls for KLF3 and SP2. **J**) Change in gene expression of *ANKRD55*/*IL6ST* (x-axis) in stimulated versus resting CD4+ T cells. **K**) Expression of *IL6ST* upon CRISPRi-mediated enhancer inhibition in primary human CD4+ T cells. * = p-value <0.05, ** = p-value <0.01, *** = p-value <0.001.

Among the top prioritised genes was also *ANKRD55* (**Figure 7A**), a gene which has not been very well described in T cell biology hitherto. *ANKRD55* is a pleiotropic gene identified in all five eGRNs, and it is regulated by an enhancer containing four AD associated genetic variants from multiple sclerosis (MS), JIA, rheumatoid arthritis (RA) and inflammatory bowel disease (IBD).^70^ Additionally, the enhancer regulating *ANKRD55* in our eGRN (chr5:56139147-56162711) was also connected to *IL6ST* (**Figure 7G**), a gene which has been associated with T cell function and disease activity in SLE.^71^ The enhancer overlaps two AD associated peaks downregulated in most ADs we analysed (**Figure 7H**), except for peak chr5:56139295-56140114, which showed a distinct upregulation in Graves’ disease. In line with this, TF activity of HIC2 (previously identified as a transcriptional repressor^72^, and linked to enhancer chr5:56139147-56162711 in our eGRN) was increased in Graves’ disease, while being decreased in psoriasis and SLE (**Figure 7I**). On the contrary, activity of SP1 (linked to the same enhancer and previously identified as a transcriptional activator^73^), was decreased in Graves’ disease and increased in psoriasis and SLE (**Figure 7G**). Moreover, we found *ANKRD55* and *IL6ST* expression to be downregulated in activated versus resting T cells (**Figure 7J**), indicating they play a role in limiting T cell activation. CRISPRi perturbation of chr5:56139147-56162711 led to an increase in *IL6ST* expression (**Figure 7K**), suggesting that this enhancer acts as a repressor. *ANKRD55* expression could not be detected in our experimental setting, in line with what was observed in activated Th1 and Th17 cells (**Figure 7J**).

These results demonstrate that our eGRNs can be used to identify enhancer-gene links which are highly relevant to T cell activation in AD, and can be functionally validated. Similarly, we have made prioritised lists of genes which are potentially mis-regulated in SLE, psoriasis, Graves’ disease and JIA using their respective disease specific eGRN (**Supplemental Table 8**). These lists can be used to identify novel genes and their upstream transcriptional/epigenomic regulators potentially important for T cell function and the pathogenesis of specific ADs.

## Discussion

The interpretation of non-coding genetic variants associated with ADs remains a significant challenge. A deeper comprehension of the role of SNPs and regulatory elements in the dysregulation of T cell activation in ADs is necessary for improved understanding of disease mechanisms and the identification of potential new therapeutic targets. To link SNP harbouring enhancers to their downstream target genes, we constructed AD specific eGRNs comprising genetic and epigenetic evidence from CD4+ T cells, that can be used to identify targets that inhibit aberrant T cell activation.

From the enhancer landscape, we observed that AD T cells are characterised by general as well as disease specific enhancer signatures that potentially regulate disease relevant biological pathways. This was also reflected on the TF activity level, where we identified disease specific changes but also some core differential TFs that had an increased activity across all diseases. The shared TFs were related to T cell activation and inflammation, common processes which are dysregulated in all autoimmune diseases. These TFs included members of the AP-1 complex (including JUN and FOS family members), which are known to be activated through T cell receptor (TCR)-mediated and co-stimulatory receptor (i.e. CD28) signalling.^74^ This shows that the shared dysregulation of T cells in AD are mainly related to general T cell activation, likely due to general antigen binding by the TCR. The disease specific TF activity signatures could be driven by specific signalling related to the nature of the antigen triggering T cell activation, independent from the TCR. Indeed in SLE we find an increased activation of IFN related TFs (in line with increased IFN signalling in these patients^26^), in Graves’ disease we find TFs from the C/EBP family which are involved in IL-6 signalling^75^ (a cytokine increased in the serum of Graves’ disease patients^76^), while in psoriasis we find KAISO and CREB family TFs that are involved in TGFβ signalling^43–45^ (which is also implicated in psoriasis pathogenesis^46^). Thus, we show that the epigenomic landscape of T cells can be used to identify TFs that are driving specific gene signatures in different ADs, and serves as a readout to extrapolate specific signalling pathways in T cells important for disease pathogenesis.

In addition to pinpointing disease-relevant signalling pathways through TF activity, we can harness the power of TF-enhancer-gene connections within our AD-specific eGRNs to uncover distinctive gene signatures associated with T helper subsets linked to specific ADs. We demonstrate that integrating eGRNs with expression data from Th cells offers invaluable insights into both the prevalence and functionality of particular Th subtypes within the general CD4+ T cell population. By assessing the correlation between disease-specific changes in H3K27ac signal and differential expression of linked genes upon Th subset stimulation, we shed light on the nuanced roles of these cells in different ADs. As an example, in the case of psoriasis, a disease where Th17 cells are considered pivotal drivers of pathogenesis^10^, we found evidence for the presence of Tregs that can differentiate into IL-17-producing cells, distinct from the classical Th17 phenotype^60^, and indicative of the dynamic interplay between these cell subsets. Importantly, it has been shown that the HDAC inhibitor Trichostatin A can inhibit the conversion of Tregs to Th17-like cells, highlighting that epigenetics play a key role in this process.^60^ Likewise, in SLE, we found a potential role for non-classical IFN-γ-producing Th17 cells in the disease pathogenesis. Over the last years, there has been an increasing interest in the therapeutic inhibition of Th17 cells in ADs. However, these cells display a very high level of plasticity and their origin and exact pathogenic function in autoimmunity is still poorly understood. Indeed, classical approaches to inhibit Th17 cells in autoimmunity, including IL-17A blockade, are thought not to be effective against “non-classical” Th17 subtypes^77^, demonstrating the need for a better understanding of the exact Th subtypes involved in AD. Our findings not only emphasise the heterogeneity of T cell responses in AD but also shed light on potential therapeutic targets, for example through the modulation of specific enhancers linked to genes involved in the pathways driving specific T cell subsets.

We also show that AD specific eGRNs are enriched in super enhancers that regulate processes related to T cell activation and inflammation. Recent work has shown that BET (Bromodomain and extra terminal) and Mediator proteins are crucial for transcriptional activity of these super enhancers^78^, and inhibition of these bromodomains are being pursued as therapeutics to control the aberrant rate of transcription of disease genes in cancer and other diseases.^79^ JQ1 is one of the well-studied BET inhibitors, and has shown to repress the expression of T cell activation related genes in memory/effector T cells in JIA.^20^ In line with this, we show that JQ1 selectively inhibits the expression of genes linked to super enhancers identified in our JIA specific eGRN. Since we also found an enrichment of super enhancers in our general AD and psoriasis eGRNs, these results could potentially be extrapolated to other ADs. Indeed, JQ1 treatment has previously been shown to alleviate general inflammation as well as a decrease in the number of CD4+IL-17A+ T cells in imiquimod-induced psoriasis mouse models^80,81^, further highlighting the therapeutic potential of BET inhibitors in the treatment of AD.

Enhancers are traditionally often linked to their nearest genes using proximity based methods, even though it is known that they can regulate their target genes over large genomic distances^82^, and autoimmune associated SNPs in non-coding regions are often affecting distal genes.^83^ Our eGRNs can be used to identify distal enhancer-target gene links that would be missed using more conventional proximity based methods. As an example, we identified a distal enhancer linked to *CTLA4* (chr2:203930706-203942262, with a distance of 60052 bp), that was also linked to *ICOS* (TSS, 0 bps) and *CD28* (222190 bp). This enhancer overlaps six SNPs (rs56324422, rs4675374, rs11571305, rs11571306, rs4335928 and rs10932031) that have previously been identified as a Celiac disease risk variants.^52,84^ Although these SNPs have mainly been associated with *ICOS* expression (the most proximal gene), we show that perturbation of the enhancer harbouring these SNPs also affects *CTLA4* and *CD28* which are more distal as predicted by our eGRN. *ICOS*, *CTLA4,* and *CD28* play crucial roles in regulating T cell responses^66^, and dysregulation of their signalling can lead to the breakdown of immune tolerance. Likewise, we also identified *IL6ST* and *ANKRD55*, two genes that we found to be regulated by an enhancer located at chr5:56139147-56162711 that overlaps a T cell specific super enhancer and harbours many AD associated SNPs. One of these SNPs, rs6859219 has previously been associated with increased *ANKRD55* transcript levels in PBMCs and CD4+ T cells.^85^ Moreover, increased *IL6ST* expression has been shown to increase Th17 differentiation (a cell type highly relevant in AD) in mouse CD4+ T cells.^86^ In our eGRN we identified HIC2 as a TF that negatively regulates the expression of *ANKRD55* and *IL6ST* through the enhancer at chr5:56139147-56162711, in line with the previously described repressor function of HIC2.^72^ SNPs in this enhancer could potentially disrupt the repressive function of HIC2, thereby leading to increased expression levels of *ANKRD55* and *IL6ST*. Indeed, we show that CRISPRi mediated perturbation of the enhancer harbouring the HIC2 binding motif leads to an increase in *IL6ST* expression in primary human CD4+ T cells. HIC2 function has not been studied in T cells or other immune cells hitherto, but our analysis points to it as a potentially important regulator of *IL6ST* and possibly also of *ANKRD55* and its enhancer in the context of AD. This demonstrates the potential of eGRNs to link AD associated enhancers harbouring SNPs to downstream genes as well as upstream regulators, and uncover new regulatory mechanisms driving T cell dysfunction in autoimmunity.

Taken together, here we demonstrate that eGRNs are powerful tools to study disease specific dysregulation of T cells. They can be used to pinpoint alterations in the regulatory landscapes that contribute to the development of ADs and provide a basis for developing new therapeutic approaches.

## STAR Methods

## KEY RESOURCES TABLE

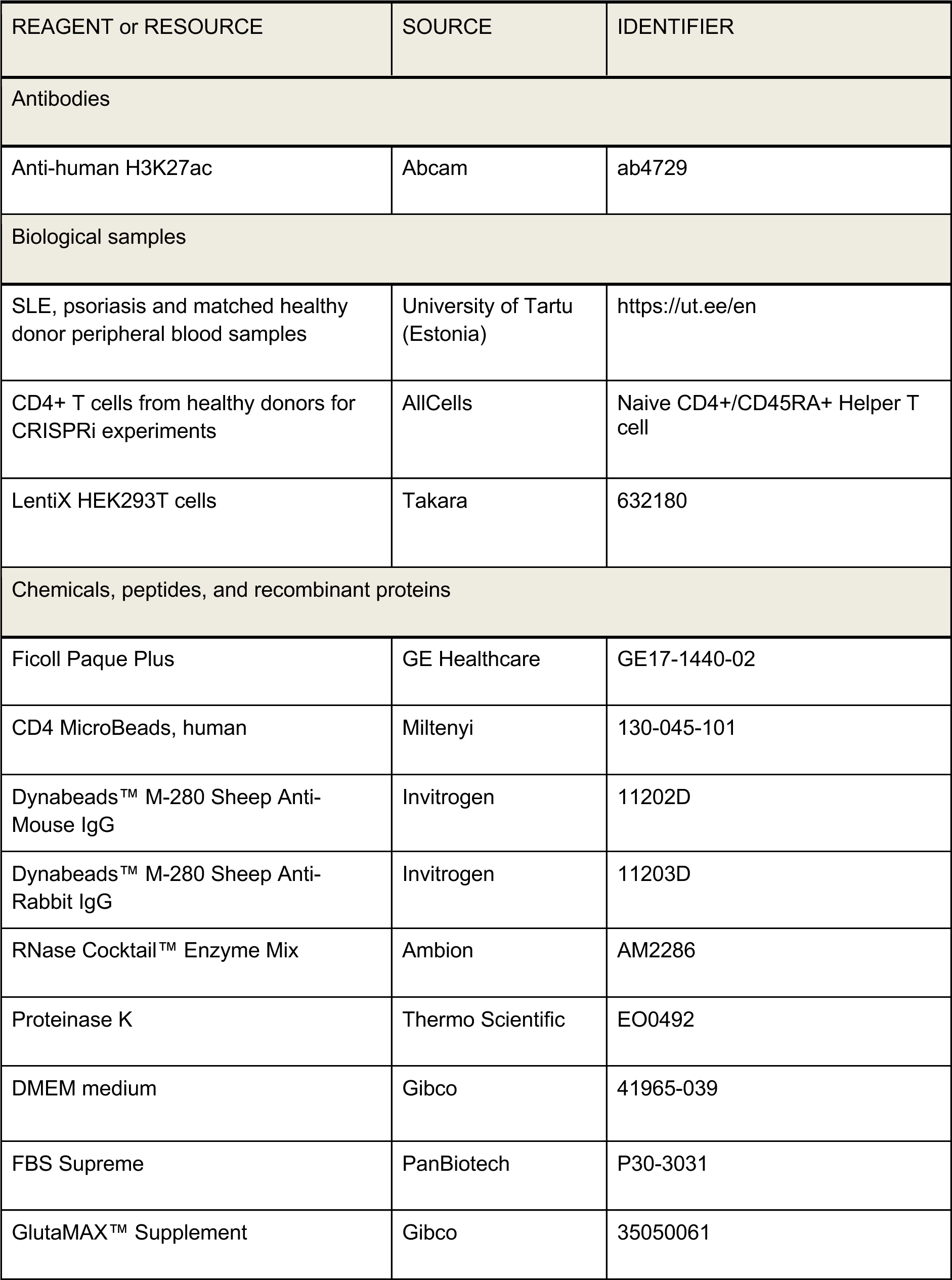
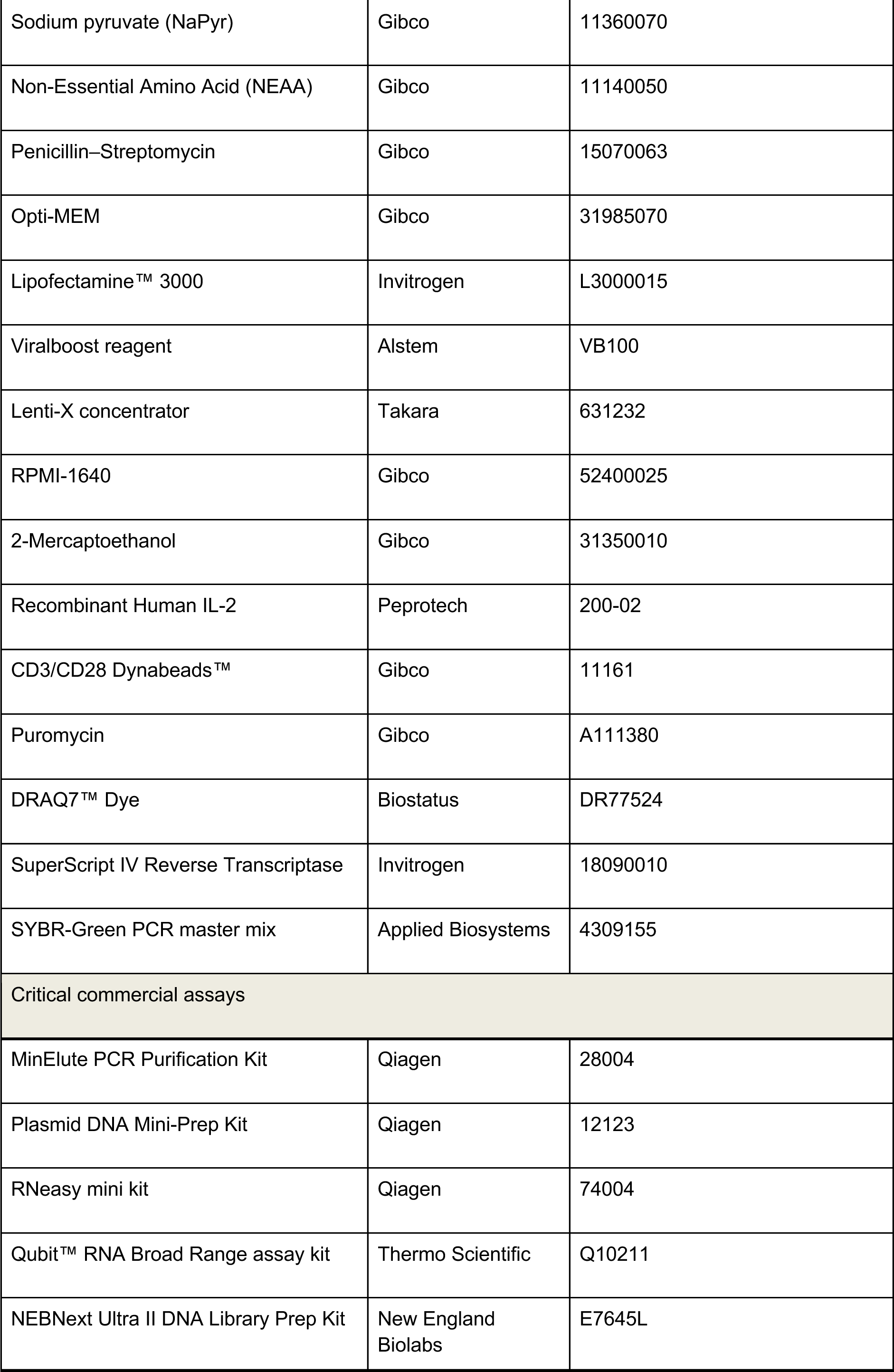
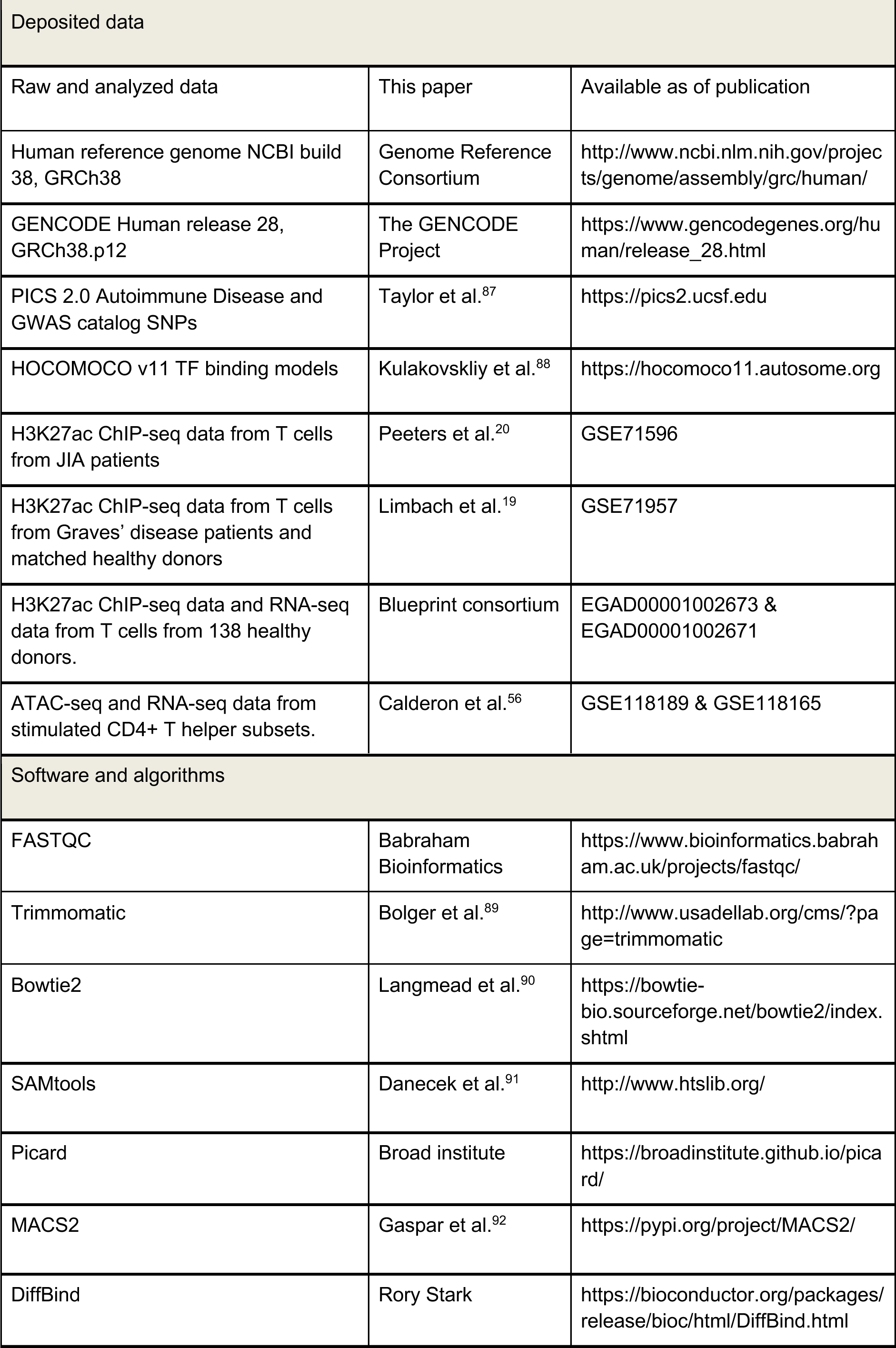
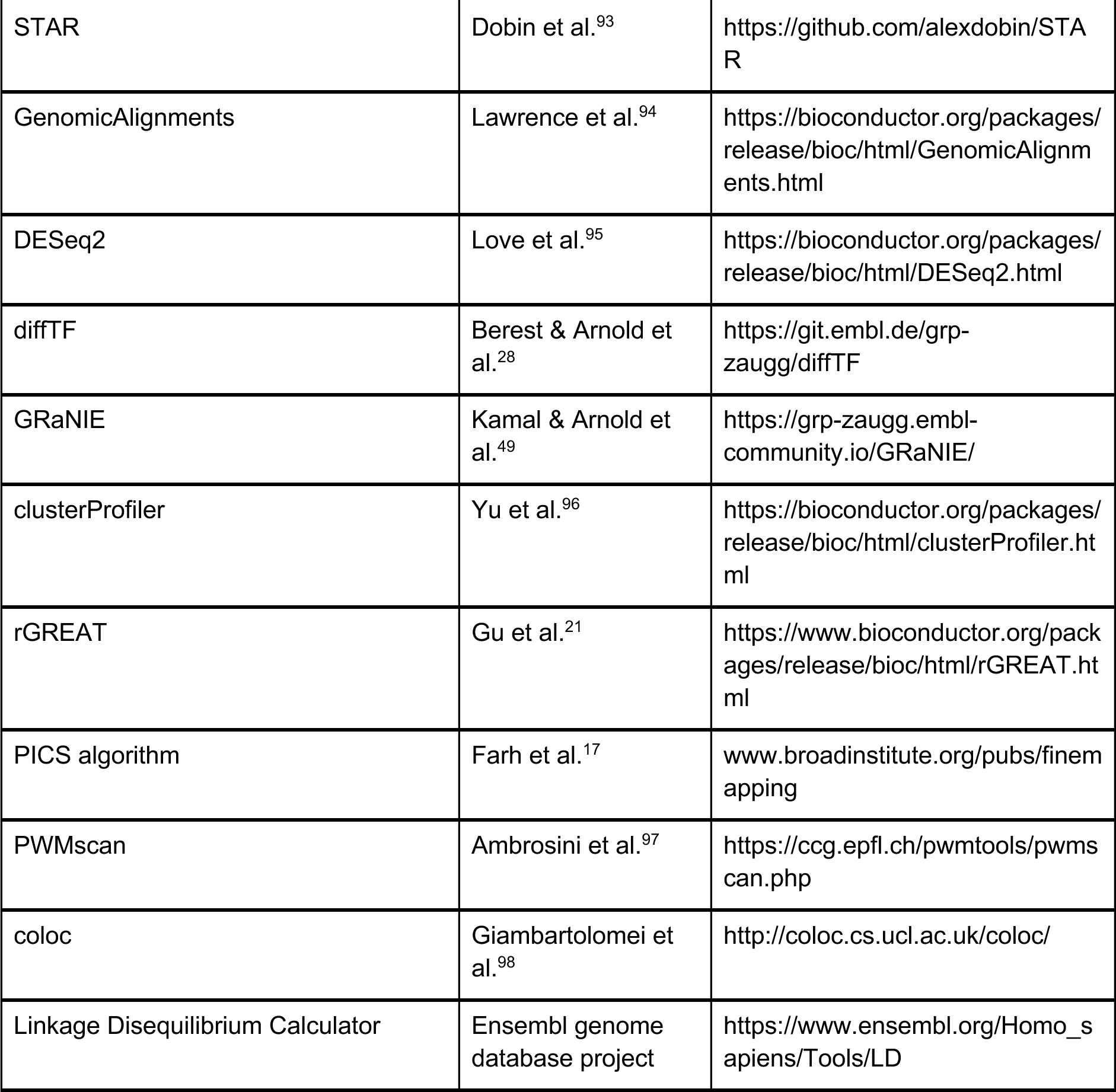

## RESOURCE AVAILABILITY

### Lead contact

Further information and requests for resources and reagents should be directed to and will be fulfilled by the lead contact, Judith Zaugg.

### Materials availability

This study did not generate new unique reagents.

### Data and code availability

De-identified patient H3K27ac ChIP-seq data will be publicly available as of the date of publication. Additionally, this paper analyzes existing, publicly available data. The accession numbers for these datasets are listed in the key resources table. All original code has been deposited at https://gitfront.io/r/user-2800625/inXS3mECwiLE/AD-Enhancer-Remodelling/ and is publicly available as of the date of publication. Any additional information required to reanalyze the data reported in this paper is available from the lead contact upon request.

### Author Contributions

Conceptualization, N.D., and J.B.Z.; Methodology, N.D., C.A., A.R.P.; Software, N.D., N.H.S., C.A., and A.R.P.; Validation, N.D., N.H.S. and D.M.; Formal analysis, N.D., and N.H.S.; Investigation, D.M., K.K., and E.K.; Resources, K.K., K.K., R.K., K.U., L.S., P.P., and J.B.Z.; Data Curation, N.D., and N.H.S.; Writing – Original Draft, N.D., N.H.S., and J.B.Z.; Writing – Review & Editing, N.D., N.H.S., K.K., D.M., C.A., A.R.P., L.S., P.P., N.N., and J.B.Z.; Visualization, N.D., and N.H.S.; Supervision, N.N., and J.B.Z.; Funding Acquisition, N.N. and J.B.Z.

### Funding

N.D., N.H.S. J.B.Z. acknowledge funding from GSK through the EMBL-GSK collaboration framework (3000032294). N.H.S., J.B.Z. acknowledge funding from the EMBL Infection Biology Transversal theme. K.K. and P.P. acknowledge funding from the Estonian Research Council (PRG377 and PRG1117) and from the Centre of Excellence for Genomics and Translational Medicine funded by the European Regional Development Fund (Project No.2014-2020.4.01.15-0012). A.R.P. has been recipient of a postdoctoral fellowship granted by Fundación Ramón Areces (2017-T2/BMD-5532). This project is co-funded by the European Union (ERC, EpiNicheAML,101044873) to J.B.Z. Views and opinions expressed are however those of the author(s) only and do not necessarily reflect those of the European Union or the European Research Council. Neither the European Union nor the granting authority can be held responsible for them.

## Acknowledgements

We thank Annique Claringbould for her feedback and discussions on integration of GWAS data in the eGRNs, and the staff at the EMBL Genomics Core Facility, and the Flow Cytometry Core Facility, for sample preparation and data generation. We are very grateful to all the patients and healthy donors who donated the samples included in this study.

## EXPERIMENTAL MODEL AND STUDY PARTICIPANT DETAILS

This study was approved by the Ethics Review Committee of Human Research of the University of Tartu, Estonia (protocols 270/T-9, 271/M-35). Informed consent was obtained from all the participants in accordance with the Declaration of Helsinki. Patient demographics are listed in **Figure 1A**.

## METHOD DETAILS

### Isolation of CD4+ T cells

30mL of peripheral blood was collected in EDTA tubes from patients with SLE (n=11) and psoriasis (n=15) and from age- and sex-matched healthy donors (n=11 and n=15 respectively) from The University of Tartu, Estonia. PBMCs were obtained by Ficoll Paque (GE Healthcare) density gradient centrifugation. CD4+ T cells were collected by magnetic isolation using CD4 MicroBeads (Miltenyi) on the AutoMACs (Miltenyi) according to the manufacturer’s instructions. For CRISPRi experiments, frozen healthy donor naive CD4+ T cells were obtained from AllCells.

### H3K27ac ChIP-seq protocol

1 million isolated CD4+ T cells were cross-linked with an 11% formaldehyde buffer, lysed and subjected to sonication using a Bioruptor sonicator UCD-200 (Diagenode) to obtain chromatin fragments of 100–500 bp. Sheared chromatin was immunoprecipitated with 2 μL of anti-H3K27ac antibody (Abcam, ab4729) and 2 μL of Dynabeads M-280 magnetic beads (Sheep Anti-Mouse IgG, Sheep Anti-Rabbit IgG, Invitrogen) using the SX-8G IP-Star Automated System (Diagenode). ChIP samples were decrosslinked at 65°C for 4 hours. After incubation, the samples were treated with 0.2 mg/mL RNaseA (RNase Cocktail Enzyme Mix, Ambion, Life Technologies) and 0.4 mg/mL Proteinase K (Thermo Scientific). DNA was purified with the MinElute PCR Purification Kit (Qiagen) according to the manufacturer’s protocol. Sequencing libraries were prepared using the NEBNext Ultra II DNA Library Prep Kit (New England Biolabs) for Illumina according to the manufacturer’s instructions. Library quality and concentration were determined using an Agilent Bioanalyzer DNA1000 chip according to the manufacturer’s instructions. After passing quality control, libraries were sequenced on an Illumina NextSeq 500 (75-bp single-end mode).

### CRISPRi in CD4+ T cells

Individual gRNAs (4 per enhancer, 2 for CD4 promoter and 2 non-targeting scrambled gRNAs, **Supplemental Table 4**), were cloned into the CROP-seq Puro-F+E plasmid, containing the optimised F+E tracrRNA sequence ^99^, using restriction ligation as previously described.^100^

Lentivirus was produced in LentiX HEK293T cells (Takara). Cells were grown to 65–75% confluency prior to transfection. Lentiviral packaging vectors pMD2.G (Addgene 12259) and psPAX2 (Addgene 12260) were combined with CROPseq-Puro-F+E plasmid (containing the sgRNAs) or dCas9-mCherry-ZIM3-KRAB plasmid (Addgene 154473) and transfected using Lipofectamine™ 3000 (Invitrogen). After 6 hours, the transfection medium was replaced with a complete medium supplemented with 1:500 Viralboost reagent (Alstem). Supernatant containing viral particles was harvested after 2 and 3 days and concentrated 100x using Lenti-X concentrator (Takara) following manufacturers instructions.

For T cell transduction, frozen healthy donor naive CD4+ T cells were thawed, cultured and activated using Human T-Activator CD3/CD28 Dynabeads™ (Gibco). The following day, cells were infected with dCas9-mCherry-ZIM3-KRAB lentivirus and the next day with gRNA lentivirus. Cells were selected for gRNA expression using Puromycin and were subsequently reactivated. After three days of reactivation, mCherry positive T cells expressing the dCas9-ZIM3-KRAB protein were sorted by fluorescence-activated cell sorting (FACS) on a BD FACSAria™ Fusion, selecting for viable cells using DRAQ7™ Dye (Biostatus). Sorted cells were immediately lysed in RLT plus buffer (Qiagen) supplemented with 2-Mercaptoethanol.

The CRISPRi effect on predicted enhancer–target genes was measured using quantitative PCR. Briefly, RNA was isolated using the RNeasy mini kit (Qiagen) according to the manufacturer’s instructions. RNA concentration was measured using the Qubit™ RNA Broad Range assay kit (Invitrogen), and reverse transcription was performed using SuperScript IV according to the manufacturer’s instructions. qPCR was performed with SYBR-Green PCR master mix (Applied Biosystems), and data was processed using the -ΔΔCt method, normalised to the housekeeping gene GAPDH. Primer sequences can be found in **Supplemental Table 9**.

## QUANTIFICATION AND STATISTICAL ANALYSIS

### ChIP-seq pipeline

Reads were trimmed using Trimmomatic v0.36 for illumina TruSeq3-SE adaptors. FASTQC was performed on samples before and after trimming and those samples which passed quality control were used for alignment. The ChIP-seq reads were then aligned to human genome assembly hg38 using Bowtie2^90^ with default parameters in “*—very-sensitive*” mode. Aligned reads mapping to ChrM, other unassembled contigs and also reads with minimum mapping quality score < 10 were filtered out using SAMtools v1.5.^91^ Additionally, duplicates were removed using Picard tool v2.17.6 using the “*MarkDuplicates”* function.

Peak calling was done per sample using MACS2^92^ v.2.0.1 using default parameters and “*--keep-dup all*”. Consensus peaks for the samples within each disease were generated with the DiffBind Bioconductor R package using the “*dba.peakset*” function with the requirement that the peaks are present in at least two samples in case of Graves’ and JIA and at least three samples in case of Psoriasis and SLE. A consensus peakset for all the diseases together (Psoriasis, SLE, Graves’ and JIA) was generated with the peaks present in at least three samples.

The raw counts for the resulting consensus peak sets were generated using the “*dba.count*” function with default parameters.

### RNA-seq pipeline

Reads were trimmed using Trimmomatic^89^ v0.36 for Illumina TruSeq3-PE-2 adaptors. FastQC was performed on samples before and after trimming and those samples which passed quality control were used for alignment. The reads were then aligned to human genome assembly hg38 using STAR^93^ v2.6.0 with default parameters. Aligned reads which were uniquely mapped and in correct pairs were retained. Gene counts were obtained using the “*summarizeOverlaps*” function from the GenomicAlignments Bioconductor R package^94^, using the gencode.v28 primary assembly annotation gtf file.

### Differential peak/gene analysis

PCA and hierarchical clustering were performed on transformed read counts (regularized log transformation) for peaks/genes in case of ChIP-seq and RNA-seq respectively. This was done as a quality control measure to identify sample similarity based on conditions and to detect any kind of batch effects prior to performing differential peak/gene analysis. Differential analysis for peaks/genes were carried out using the DESeq2 Bioconductor R package^95^ with default settings and using the following generalized linear model: “*∼ Age + Batch + Condition*”. Peaks and genes with an adjusted p-value threshold <= 0.05 were considered significant.

### Differential TF activity

Differential TF activity between autoimmune patients and controls was estimated using the diffTF tool^28^ in analytical mode with default parameters and using HOCOMOCOv11 motifs (as part of the DiffTF workflow). TFs with an adjusted p-value <=0.05 were considered significantly differential.

### GWAS-SNPs for autoimmune diseases

The summary statistics for genome-wide association studies for ADs including psoriasis^101^, SLE^102^, IBD^103^, RA^104^, celiac disease^84^, and asthma^105^ were downloaded from NHGRI-EBI GWAS-catalog^52^ for the hg38 genome build.

SNPs with p-value < 1e-5 and those mapping not in the MHC region (ch6:20,000,000 – 40,000,000) were selected. These selected GWAS-SNPs together with their linkage disequilibrium (LD) variants, obtained for the hg38 genome build, were used in this analysis. These LD variants were identified using a window of 200kbp and R squared value equal or greater than 0.8 (R2 >= 0.8) around selected GWAS-SNPs used as lead SNP using the LD calculator from https://www.ensembl.org/Homo_sapiens/Tools/LD.

### Colocalization analysis of hQTLs and GWAS-SNPs

We used coloc v2.3.1^98^ to perform colocalization between hQTLs from naive T cells and GWAS-SNPs belonging to several ADs using their summary statistics as stated above. The hQTL data was obtained from blueprint consortium.

Colocalisation was performed on 400kb regions centered on significant hQTL peaks (p-value < 0.05). In this region, we ran colocalization tests if the number of shared hQTLs and non HLA-GWAS SNPs were greater than 10 (shared SNPs > 10 in 400 kb region). We considered a true colocalization signal, if the probability of association between GWAS-SNP and hQTL due to a shared causal variant given as PP.H4 by coloc, was greater than or equal to 0.8 (PP.H4 >= 0.8).

### Generation of naive CD4 T-cell eGRN

We obtained paired RNA-seq and H3K27ac ChIP-seq data from naive CD4+ T cells from 138 healthy donors from the Blueprint consortium to generate eGRN. Both datasets were downloaded from the EGA archive (EGAD00001002671, EGAD00001002673) and processed using ChIP-seq and RNA-seq analysis pipeline as described above. It is based on covariation between enhancer signal and RNA-expression across individuals, adapted from the workflow described in^49^. The eGRN generation consist of two main steps:

1. Generation of TF-enhancer links

We obtained putative TF binding sites using PWMscan^97^ for all the TFs for which the motif is available from HOCOMOCO v.11. ^88^ Next, we identified the TF binding sites which overlapped the consensus H3K27ac ChIP-Seq peak set (generated as described above). We then generated a correlation matrix between TFs (columns) and H3K27ac peaks (rows). Each cell in the matrix is a Pearson correlation coefficient between TF expression and H3K27ac signal across all 138 individuals for the corresponding TF-peak pair. We then calculate FDR for each TF-enhancer pair as described in ^51^. We used an FDR threshold of 20% for all TFs to define significant TF-enhancer links.

2. Generation of enhancer-gene links

Enhancers were linked to all the genes within a distance of +/- 250kb TSS and which show positive and significant Pearson correlation coefficient between H3K27ac signal and gene expression across all 138 individuals with an adjusted P-value cutoff of 0.05.

Next, we combined the two steps above which resulted in the generation of a tripartite eGRN containing TF-enhancer-gene links.

Additionally, we include SNP-enhancer-gene links by overlapping known AD SNPs obtained from NHGRI-EBI GWAS-catalogue as described above with the enhancers included in the consensus peak set and correlating these enhancers to putative downstream target genes as described in step 2. Similarly as above, we also include Differentially acetylated-enhancer-gene links by overlapping differentially acetylated enhancers (AD patient vs controls) with consensus enhancer peaks.

### Generation and pruning of AD specific eGRNs

The autoimmune eGRNs for each disease (Graves’ disease, SLE, JIA and psoriasis) were generated separately. Each of these disease-specific eGRNs involved filtering the naive CD4+ T cell eGRN to comprise only nodes that are linked to at least two different or two of the same types of disease specific molecular evidence. These molecular evidences include (i) differentially active TFs, in which case we would keep their regulatory elements and target genes, (ii) differentially acetylated enhancers, in which case we would keep their upstream linked TF and their downstream linked target gene. In addition, we keep enhancers and their linked TF and target genes if (iii) the enhancer overlaps with a GWAS-SNP of the corresponding disease, or (iv) if enhancer overlaps with with naive T cells peaks associated with significant colocalized histone quantitative trait loci (hQTLs) -AD GWAS SNPs

We applied additional criteria to this filtered network. Specifically, a gene is kept if any of the following is true:

a. It is connected to more than one differentially active TF with their regulatory activities going in the same direction (either upregulation or downregulation).
b. It is connected to more than one differential H3K27ac peak with their activity going in the same direction (either upregulation or downregulation)
c. It is connected to a differentially active TF AND a differentially active peak with their regulatory activities in the same direction
d. It is connected to a regulatory element with overlapping GWAS-SNPs/colocalized signal

### Generation and pruning of the general AD eGRN

The general AD eGRN was generated by filtering the naive CD4+ T cell eGRN in a similar way as described above but with nodes that are linked to any molecular disease evidence (differentially active TFs, differentially acetylated enhancers, GWAS-SNPs, (hQTLs) -AD GWAS SNPs peaks) coming from at least two diseases. Specifically, a gene is kept if any of the following are true:

1. It is connected to at least 2 TFs that is shared between at least 3 diseases (SLE, JIA, Psoriasis and Grave’s disease) AND that go in the same direction within at least one of the shared disease / 1 TF that is shared between at least 3 diseases (SLE, JIA, Psoriasis and Grave’s disease) AND that go in the same direction across the shared diseases
2. It is connected to at least two differential peaks that are shared between at least two diseases (SLE, JIA, Psoriasis and Grave’s disease) AND that go in the same direction across the shared diseases
3. It is connected to a peak that overlaps/is colocalized with a GWAS-SNPs associated with any one of ADs like Graves’ disease, Psoriasis, SLE, JIA, Celiac, RA, IBD and Asthma AND either a differentially active TF or peak in any of the disease covering at least two

### Prioritisation of genes from autoimmune disease networks

The genes from all AD eGRNs were prioritised first based on i) number of types of molecular disease evidence followed by ii) total number of molecular evidences. Genes are given the same priority if they have the same type and total number of molecular evidences.

For the identification of the top shared genes among different AD eGRNs, they were first selected if they were shared between 3 or more AD eGRNs. Next, they were prioritised based on following factors in the sequential order i) no. of diseases ii) number of types of molecular disease evidence and iii) total number of molecular evidences. Additionally, we have used three metrics to add more information/external validation to this prioritised list i) regulation by a fine mapped autoimmune disease variant ii) significant change in gene expression upon T cell stimulation iii) annotation from known association to autoimmune condition.

### Publicly available epigenetic data for Graves and JIA

H3K27ac ChIP-seq data from PBMC derived CD4+ T cells belonging from healthy controls and Graves’ disease patients was obtained from the NCBI gene expression omnibus using accession code GSE71957.^19^ H3K27ac ChIP-seq data for CD4+ T-cell from peripheral blood and synovial fluid from JIA patients and peripheral blood from three healthy donors was obtained using the accession code GSE71596.^20^ All samples were processed using the ChIP-seq pipeline as discussed above and used for further analysis in our study.

### Gene Ontology (GO) and GREAT enrichment analysis

The GO enrichment analysis for genes present in disease-specific and general AD networks were performed using the “*enrichGO*” function from clusterProfiler package Bioconductor R package^96^ for biological processes. All the genes in the naive T cell network were used as the background for the enrichment. Terms with an adjusted p-value cutoff <= 0.2 were considered significant.

The nearby genes in different clusters of histone acetylation signal in autoimmune disease were annotated and an enrichment analysis using GO ontology for biological process was performed using rGREAT Bioconductor R package^21^ under default settings with hg38 genome assembly. The consensus H3K27ac peak set which was used for generating clusters was used as background for calculating the enrichment. Terms with an adjusted p-value cutoff <= 0.01 were considered significant.

### Drug target enrichment

A combined list of drug targets for autoimmune diseases under consideration including psoriasis, SLE, Graves’ disease, JIA, Celiac disease, IBD, and asthma was generated. The drug targets were downloaded separately for each disease from the open targets platform https://www.targetvalidation.org/. The genes in the AD eGRNs (general plus disease-specific networks together) were tested for drug target enrichment against all the genes present in the naive CD4+ T cell network as a background using the “*fisher.test*” base function in R. Enrichments with Fisher’s exact test p-value <=0.05 were considered significant.

### Fine mapped GWAS-SNPs

Fine-mapped GWAS variants for ADs were generated using the probabilistic identification of causal SNPs (PICS) algorithm.^17^ AD variants were downloaded from i) a previously published list ^17^ and lifted to the hg38 genome build and ii) the PICS data portal under the filename “PICS2-GWAScat-2020-05-22.txt.gz” from https://pics2.ucsf.edu for the hg38 genome build. PICS probability of >50% was used to filter the variants. We identified target genes by overlapping these variants with the peaks from the naive T cell eGRN and then used the peak-gene link from this eGRN to assign the target genes.

### Stimulation landscape in CD4+T cells

ATAC-seq and RNA-seq data for CD4+ T cell subtypes belonging to Tfh, Th17, Treg, Th1 and Th2 for resting and stimulated conditions were obtained from the NCBI gene expression omnibus using accession codes GSE118189 and GSE118165 respectively.^56^ These samples were then processed using ATAC-seq pipeline and RNA-seq pipeline followed by differential peak/gene analysis between stimulation and resting conditions using DESeq2^95^ as described above.

