## Supplemental Figure for "Integration of genetic and epigenetic data pinpoints autoimmune specific remodelling of enhancer landscape in CD4+ T cells"

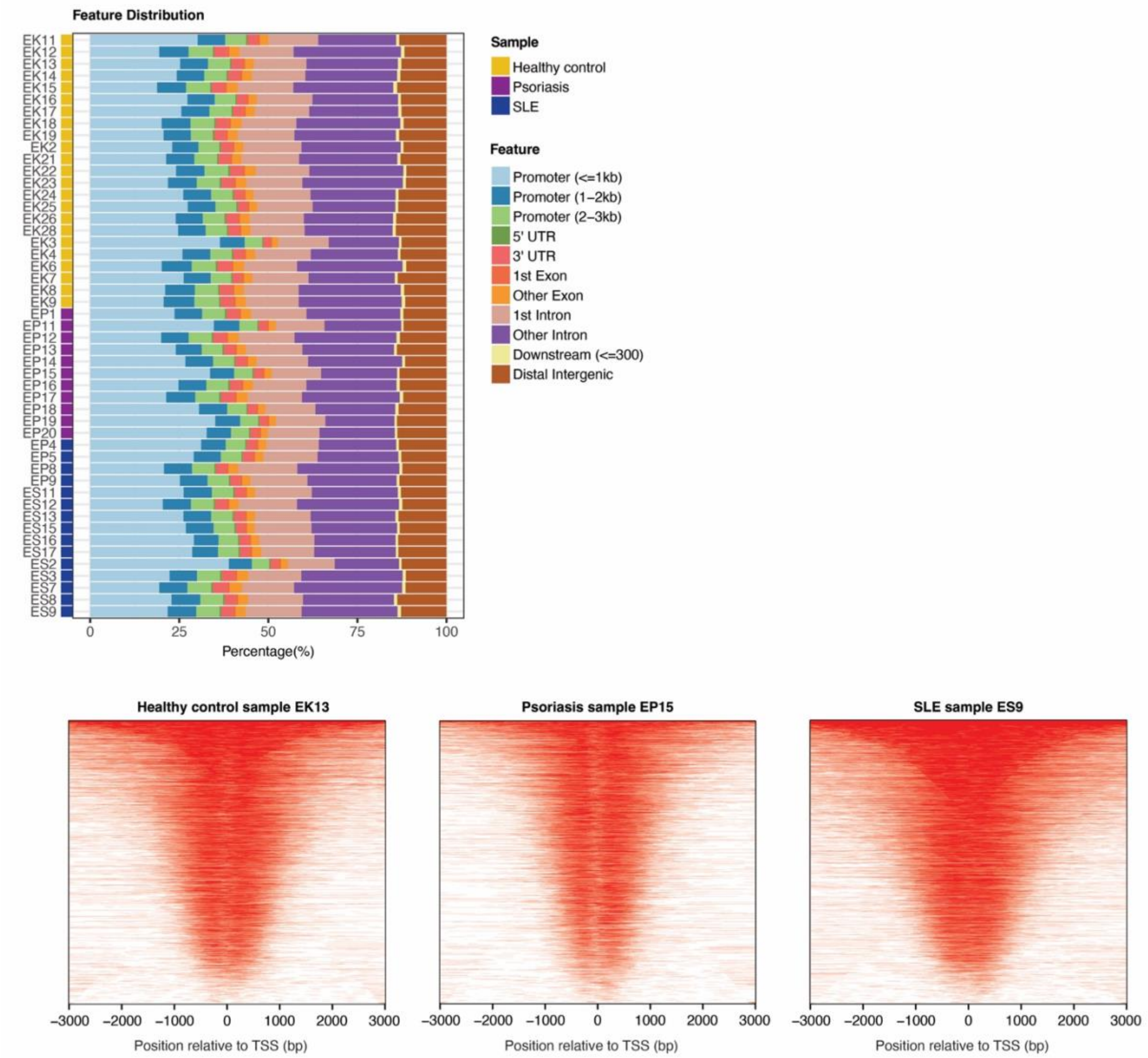


**Supplemental Figure 1. Quality control of H3K27ac ChIPseq data.** Top: Feature distribution plot of peaks across genomic features for individual samples included in the psoriasis and SLE cohorts. Bottom: tagHeatmaps visualizing the read count frequency relative to the transcription start site (TSS) for representative samples from healthy donors, psoriasis and SLE.

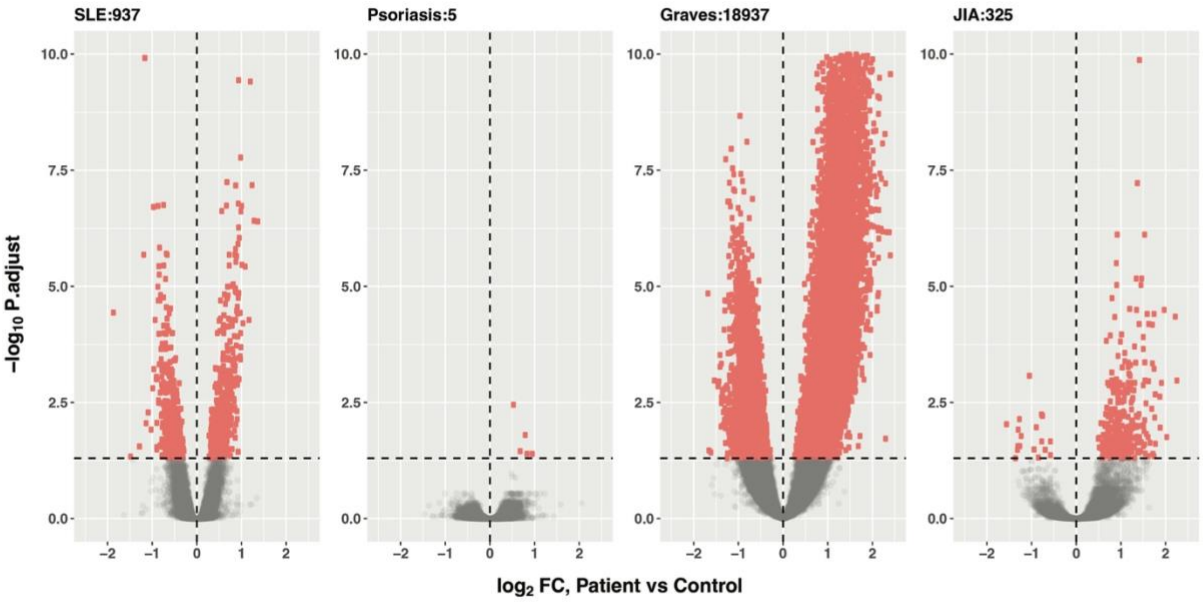


**Supplemental Figure 2. Differential peaks between AD cohorts and healthy donors.** Volcano plots highlighting differential peaks between different ADs (SLE, psoriasis, Graves' disease and JIA) and their matched healthy donors. Red dots represent significantly differential peaks (adjusted Pvalue <= 0.05). For each comparison, the number of differential peaks are indicated in the titles.

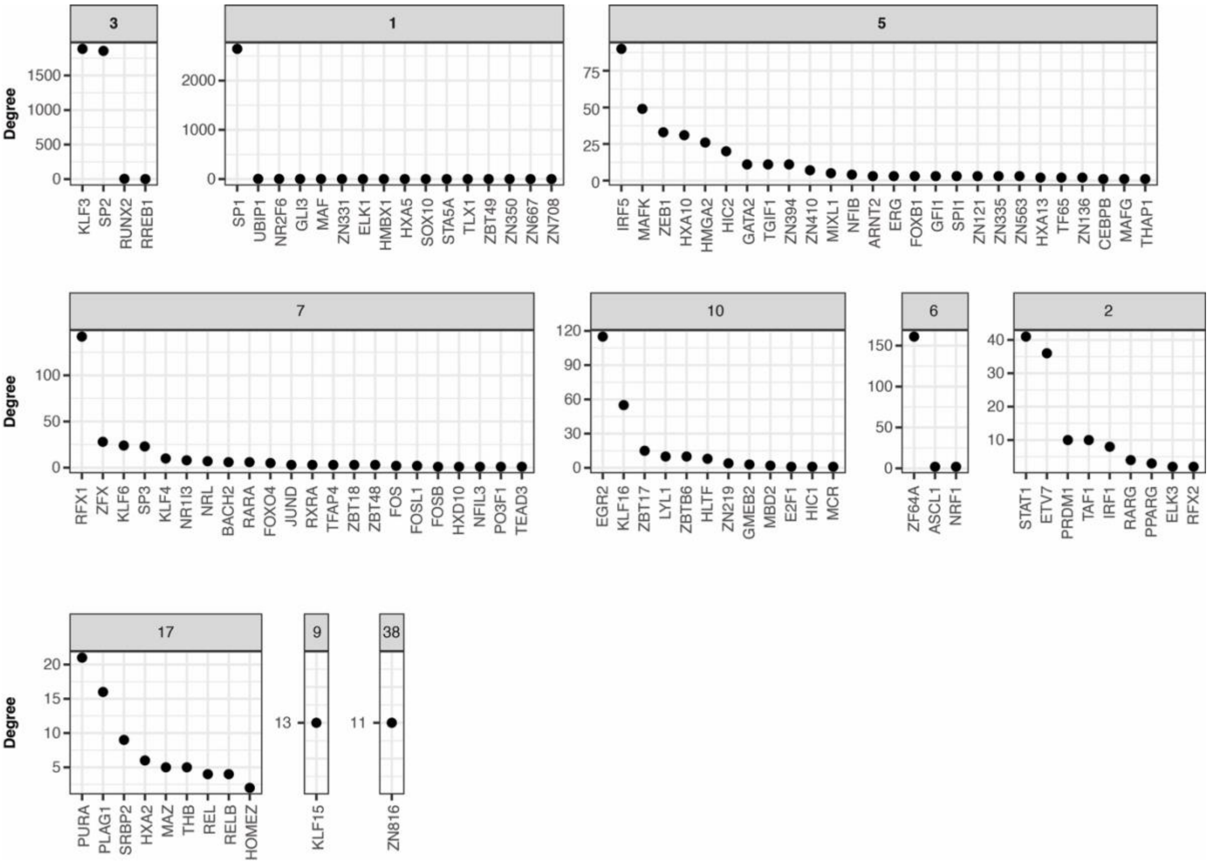


**Supplemental Figure 3. Distribution of transcription factor connections across communities.** Degree (x-axis) of transcription factors (y-axis) across communities (facet titles) identified in the naive T cell eGRN.

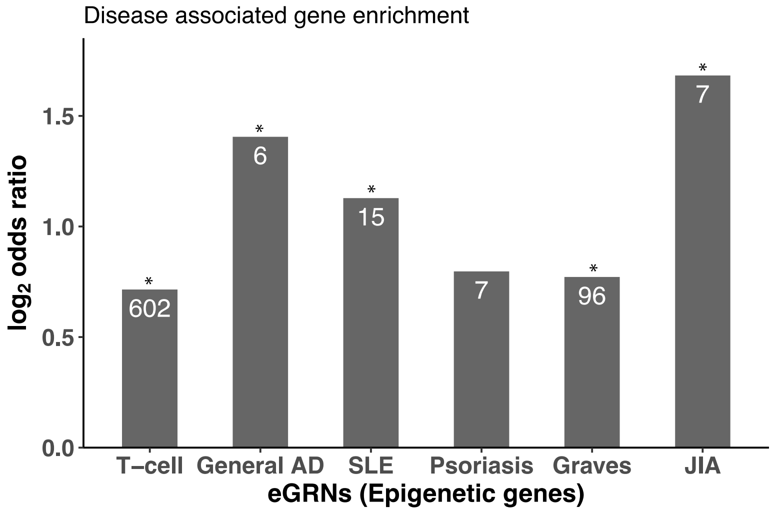


**Supplemental Figure 4. Enrichment of genes from the disease specific eGRNs with disease associated gene.** Only genes without genetic evidence were considered. Number of genes are indicated in the bars (*= Fisher's exact test p-value <0.05).

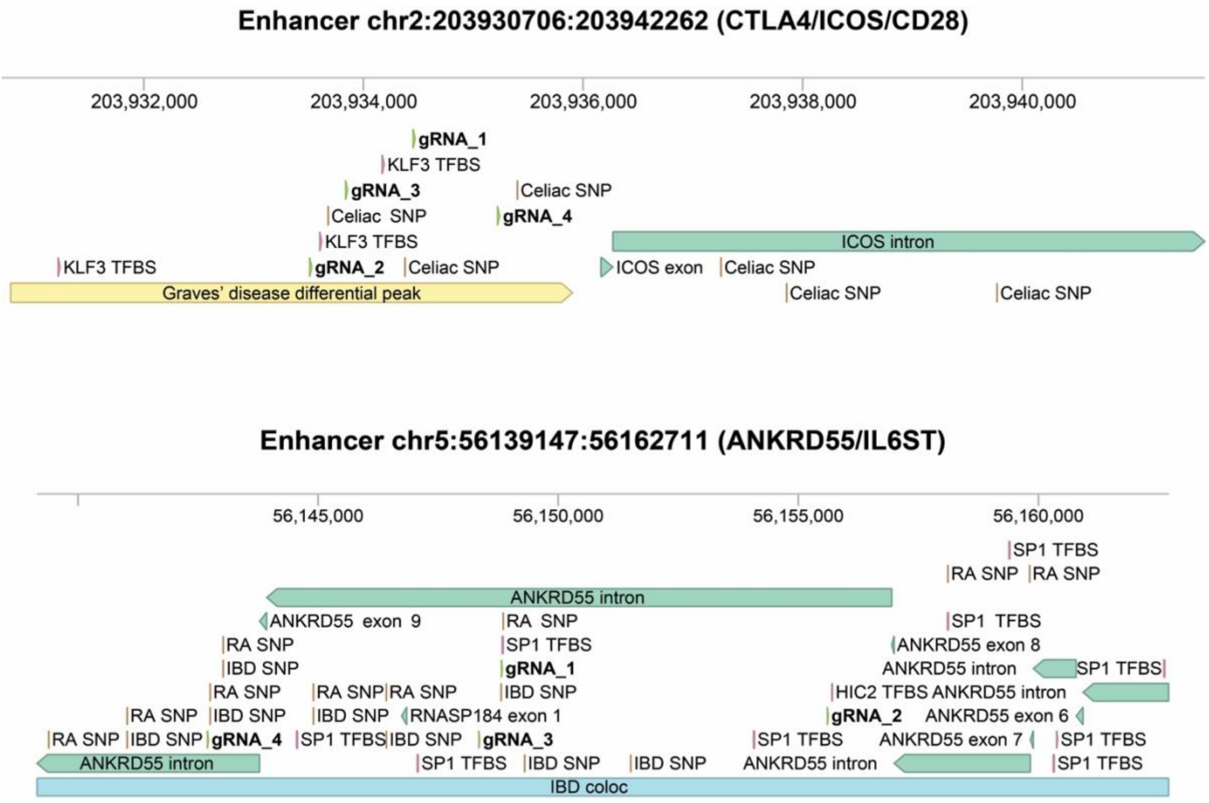


**Supplemental Figure 5. Single-guide RNAs (sgRNAs) design for CRISPRi experiments.** Genomic loci for selected enhancers linked to CTLA4/ICOS/CD28 (top) and ANKRD55/IL6ST (bottom). Position of sgRNAs (4 per enhancer) are highlighted in bold. TFBS = transcription factor binding site; SNP = single nucleotide polymorphism; coloc = colocalization of hQTLs and GWAS-SNPs.
